# Lysosome-related organelles contain an expansion compartment that mediates delivery of zinc transporters to promote homeostasis

**DOI:** 10.1101/2021.09.18.460850

**Authors:** Adelita D. Mendoza, Nicholas Dietrich, Chieh-Hsiang Tan, Daniel Herrera, Jennysue Kasiah, Zachary Payne, Daniel L. Schneider, Kerry Kornfeld

## Abstract

Lysosome-related organelles play evolutionarily conserved roles in zinc storage, but mechanisms that control zinc flow in and out are not well understood. In *C. elegans* intestinal cells, the CDF-2 transporter stores zinc in these organelles during excess. Here we identify ZIPT-2.3 as the transporter that releases zinc during deficiency. The expression levels of CDF-2 and ZIPT-2.3 are reciprocally regulated in zinc excess and deficiency, establishing a fundamental mechanism of homeostasis. Super-resolution microscopy demonstrated these organelles are composed of a spherical acidified compartment and a hemispherical expansion compartment. The expansion compartment inflates during zinc excess and deficiency by vesicle fusion delivering zinc transporters. These results identify an unexpected structural feature of lysosome-related organelles that facilitates rapid transitions in the composition of zinc transporters to mediate homeostasis.

---

Zinc plays essential roles in the structure and function of many proteins, including a predicted ten percent of the human proteome (1, 2). Zinc deficiency and excess are both toxic, and organisms require mechanisms to obtain, store, and mobilize zinc to control the amount in each cell (3, 4). Two families of eukaryotic zinc transport proteins, the Zrt, Irt-like Proteins (ZIP) and the Cation Diffusion Facilitator proteins (CDF), play critical roles in zinc homeostasis. ZIP proteins increase zinc levels in the cytosol by transporting zinc into the cytosol from the extracellular space or intracellular stores, whereas CDF proteins lower zinc levels in the cytosol by transporting zinc from the cytosol into the extracellular space or intracellular organelles (5–7). In many organisms, lysosome-related organelles have emerged as a site of zinc storage, including the yeast vacuole (8, 9), the acidocalcisome in algae (10), the zincosome in mammals (11), and gut granules in *C. elegans* (12). Critical questions about zinc storage include, how do organisms sense zinc excess and promote storage in lysosome-related organelles, and how do organisms sense zinc deficiency and promote mobilization? Here we identify ZIPT-2.3 as the *C. elegans* transporter that mobilizes stored zinc. ZIPT-2.3 and CDF-2, the transporter that stores zinc, are reciprocally regulated in zinc excess and deficiency, establishing a fundamental mechanism of homeostasis. Super-resolution microscopy revealed that lysosome-related organelles contain two compartments: an acidifed compartment facilitates degradation of macromolecules and an expansion compartment facilitates transitions in the levels of zinc transporters to mediate homeostasis.

## ZIPT-2.3 is a zinc transporter localized to lysosome-related organelles in intestinal cells

Fourteen *C. elegans* genes encode members of the ZIP protein family, named *zipt* genes (13, 14). Three of these genes, *zipt-2.1*, *zipt-2.3* and *zipt-7.1*, contain low zinc activation (LZA) enhancers in their promoter regions and are transcriptionally activated during zinc deficiency, suggesting these genes play roles in zinc homeostasis (13). To further explore regulation, we analyzed transcript levels of *zipt* genes in zinc excess. *zipt-2.3* mRNA levels were significantly lower in excess zinc conditions. This is a specific regulatory response, since 13 other *zipt* genes did not display significant regulation (Fig. S1i). Based on regulatory control in zinc deficiency and excess, we focused on *zipt-2.3*, which includes five exons and generates an 1108 nucleotide mRNA (Fig. 1a). The ZIPT-2.3 protein is highly similar to *Homo sapiens* ZIP2, *Drosophila melanogaster* dZip1(15), and *Danio rerio Dr*ZIP1 (16), suggesting these genes derived from a common ancestral gene (Fig. S2b).

**Figure 1.**
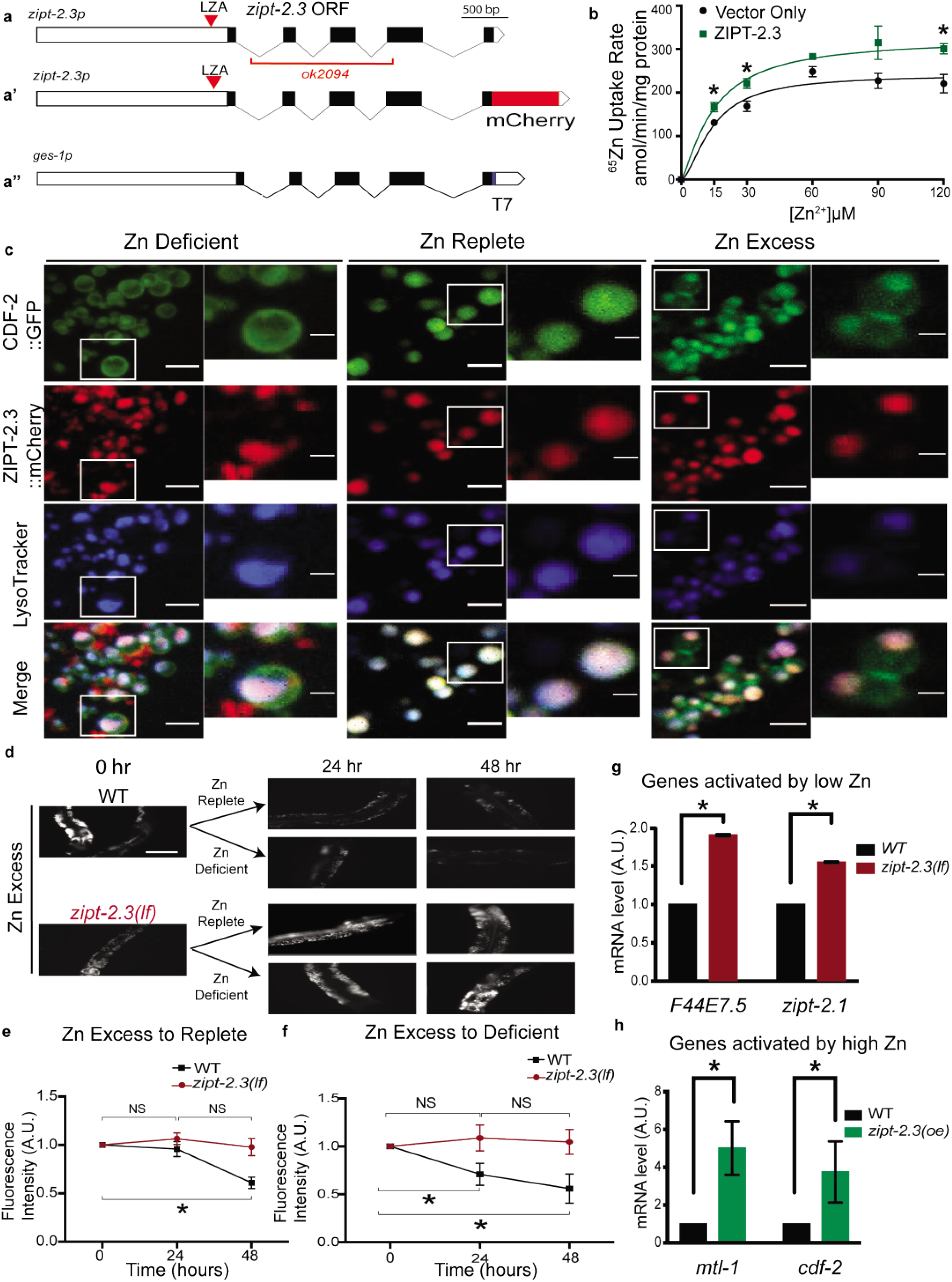
ZIPT-2.3 transports zinc from the lumen of gut granules to the cytoplasm. a-a’’) Diagrams show a portion of the plasmids in transgenic strains that express ZIPT-2.3::mCherry (a’, *amEx348* or *amEx191)* and that overexpress *zipt-2.3* in intestinal cells (a’’, *zipt-2.3(oe), amEx350*). White boxes represent promoter and 3’ UTR regions, black boxes and lines represent ZIPT-2.3 coding regions and introns, red and blue represent mCherry and T7 coding regions, the red line indicates the extent of the *ok2094* deletion mutation, and red triangles represent the LZA enhancer. b) Human HEK293T cells expressing ZIPT-2.3 or a vector control were incubated with varying concentrations of zinc containing a fixed fraction of radioactive ^65^Zn. The rate of zinc uptake was determined by measuring radioactivity that accumulated in the cells. Values are mean and SE (N= 4 replicates) (*p<0.05). c) Transgenic animals expressing CDF-2::GFP (green) and ZIPT-2.3::mCherry (red) were cultured with LysoTracker Blue in zinc replete, excess, or deficient conditions for 16 hours and visualized with confocal microscopy. Scale bars: 5 μm in larger image, and 1μm in smaller inset indicated by white box. (d-f) Wild type and *zipt-2.3(ok2094)* animals were cultured with 200μM supplemental zinc to promote zinc storage and FluoZin-3 AM to visualize labile zinc in gut granules in intestinal cells. Animals were transferred to zinc replete or zinc deficient medium (200μM TPEN), and FluoZin-3 fluorescence was analyzed by microscopy after 24 and 48 hours. (d) Representative fluorescence images show a portion of the intestine; white displays FluoZin-3 fluorescence. (e,f) Quantification of fluorescence intensity: the value at time 0 was set to 1.0 arbitrary units (AU), and other values were normalized. Values are the average of three biological replicates +/- S.D (*P<0.05). (g,h) Populations of mixed-stage, wild-type, *zipt-2.3(ok2094), or amEx350* [*zipt-2.3(oe)]* animals were cultured in standard zinc replete conditions. RNA was analyzed by qPCR. The value for WT was set to 1.0 arbitrary units (AU) for each gene, and mutant values were normalized. Values are the average of 3 biological replicates and the standard deviation (*P<0.05).

Many ZIP proteins transport zinc, but some have been shown to transport iron or other metals (17). To directly test the ability of ZIPT-2.3 to transport zinc, we expressed ZIPT-2.3 by transient transfection in human embryonic kidney cells (HEK293T) and used radioactive zinc to determine the rate of zinc uptake. Compared to vector-only control cells, cells expressing ZIPT-2.3 displayed a significant increase in the rate of zinc uptake (Fig. 1b). Thus, ZIPT-2.3 was sufficient to promote zinc uptake, consistent with the model that ZIPT-2.3 is a physiological zinc transporter (Fig. S2c).

To determine the localization of ZIPT-2.3, we generated transgenic animals expressing the *zipt-2.3* promoter and coding region fused to the coding region of mCherry (Fig. 1a’). This ZIPT-2.3::mCherry protein appears to be functional, since it rescued the *zipt-2.3(lf)* phenotype of impaired growth in zinc deficient conditions (Fig. S2d). When imaged with spinning disk confocal microscopy, transgenic animals cultured in zinc replete medium displayed a punctate pattern of expression in intestinal cells, suggestive of localization to gut granules (Fig. 1c); Chapman *et al.* (2019) reported a similar localization pattern and identified a role for *zipt-2.3* in germline apoptosis (18). To characterize this punctate pattern, we used LysoTracker, a dye that stains acidified lysosomes. ZIPT-2.3::mCherry colocalized with LysoTracker, indicating that ZIPT-2.3 localizes to the membrane of gut granules in intestinal cells (Fig. 1c).

Intestinal gut granules are the major site of zinc storage when worms are cultured with excess zinc, and the CDF-2 transporter localizes to these organelles and promotes zinc storage (12, 19). To determine if ZIPT-2.3 and CDF-2 localize to the same organelles, we generated transgenic animals that express CDF-2::GFP and ZIPT-2.3::mCherry (Fig. S2a). In zinc-replete culture medium, the gut granules appear spherical, and CDF-2::GFP and ZIPT-2.3::mCherry display complete colocalization (Fig. 1c middle). When animals are exposed to excess zinc, many gut granules display a bilobed morphology; CDF-2::GFP is localized to the membrane of both lobes, whereas LysoTracker is only localized to one lobe (12). Interestingly, ZIPT-2.3::mCherry was only localized to the LysoTracker positive lobe (Fig. 1c right). Thus, CDF-2 and ZIPT-2.3 colocalize partially during zinc excess, with both proteins on the LysoTracker positive lobe and only CDF-2 on the LysoTracker negative lobe. To examine zinc deficient conditions, we cultured animals with the zinc chelator N,N,N′,N′-Tetrakis(2-pyridylmethyl)ethylenediamine (TPEN). In zinc deficient conditions, CDF-2 and ZIPT-2.3 displayed partial colocalization. Gut granules displayed a LysoTracker positive lobe that contains both CDF-2 and ZIPT-2.3 and a small LysoTracker negative lobe that contains CDF-2 but not ZIPT-2.3 (Fig. 1c left). Thus, gut granules in intestinal cells are spherical in zinc replete conditions and remodeled during zinc excess and deficiency; in both zinc extremes there is a LysoTracker positive lobe that contains CDF-2 and ZIPT-2.3, and a LysoTracker negative lobe that contains CDF-2 but not ZIPT-2.3.

## Genetic analysis demonstrates that *zipt-2.3* promotes mobilization of zinc from gut granules and influences cytosolic levels of zinc

Based on the localization and transport activity of ZIPT-2.3, we predicted that a *zipt-2.3* loss-of-function *(lf)* mutant would be defective in mobilizing zinc stored in gut granules. To test this prediction, we analyzed the *ok2094* mutation that removes 1561 base pairs of *zipt-2.3*, including all of exons 2 and 3 and part of exon 4 (Fig. 1a). These exons encode highly conserved regions of the protein, suggesting *zipt-2.3(ok2094)* is a strong loss-of-function or null allele. We used an established assay based on the localization to gut granules of the zinc dye FluoZin-3 AM (12). Wild-type and *zipt-2.3(lf)* animals cultured with supplemental zinc and FluoZin-3 AM for 16 hours displayed strong fluorescence in gut granules, indicating robust zinc storage (Figure 1d). When shifted to zinc replete or deficient conditions, wild-type animals displayed a significant decrease in fluorescence after 24 or 48 hours, indicating stored zinc is mobilized from the gut granules. By contrast, *zipt-2.3(lf)* animals did not display reduced fluorescence, indicating a failure to mobilize stored zinc (Fig. 1d-f). Thus, ZIPT-2.3 was necessary to mobilize zinc from the gut granules.

If *zipt-2.3* releases stored zinc, then we predict that *zipt-2.3(lf)* mutants would have lower levels of cytosolic zinc compared to wild type. To test this prediction, we analyzed the expression level of zinc-regulated genes as a surrogate marker for cytosolic zinc levels. *zipt-2.1* and *F44E7.5* are activated by zinc deficient conditions (13), and the mRNA levels of both genes were increased significantly in *zipt-2.3(lf)* mutant animals compared to wild type, suggesting cytosolic zinc levels are decreased in these mutant animals (Fig. 1g). To determine if *zipt-2.3* is sufficient to increase cytosolic zinc levels, we overexpressed *zipt-2.3* in intestinal cells. We generated a transgenic strain containing multiple copies of *zipt-2.3* controlled by the *ges-1* promoter (*zipt-2.3(oe)*) that is constitutively expressed in intestinal cells (Fig. 1a’’). *mtl-1* and *cdf-2* are activated by zinc excess conditions (19, 20), and the mRNA levels of both genes were increased significantly in *zipt-2.3(oe)* mutant animals compared to wild type, suggesting cytoplasmic zinc levels are increased in these mutant animals (Fig. 1h). Thus, *zipt-2.3* was necessary to release stored zinc and maintain normal levels of cytosolic zinc and sufficient to increase cytosolic zinc levels.

## *zipt-2.3* promotes organismal zinc homeostasis

To investigate the function of *zipt-2.3* in organismal zinc homeostasis, we analyzed the growth rate of wild type and *zipt-2.3(lf)* animals in zinc deficient conditions. Synchronized L1 stage animals were cultured for 72 hours, and length was determined as a quantitative measure of growth (12). In zinc replete medium, wild-type and *zipt-2.3(lf)* animals grew and developed to similar sized adults. In zinc deficient conditions, wild-type animals displayed a slight, dose dependent growth inhibition. By contrast, *zipt-2.3(lf)* mutants displayed severe growth defects, indicating hypersensitivity to zinc deficient conditions (Fig. 2a). Mutations of six other *zipt* genes did not cause hypersensitivity, suggesting this phenotype is specific for loss of *zipt-2.3* (Fig. S1c-h). Furthermore, the *zipt-2.3(lf)* phenotype was specific for zinc deficiency, since these mutant animals did not display hypersensitivity to the iron chelator 2,2-bipyridyl or the manganese chelator diaminocyclohexanetetraacetic acid (DCTA) (Fig. 2b, S1b). Furthermore, *zipt-2.3(oe)* animals displayed hypersensitivity to excess zinc compared to the wild type (Fig. 2c). Thus, *zipt-2.3* was necessary for growth and development in zinc deficient conditions and sufficient to cause hypersensitivity to high zinc toxicity.

**Figure 2.**
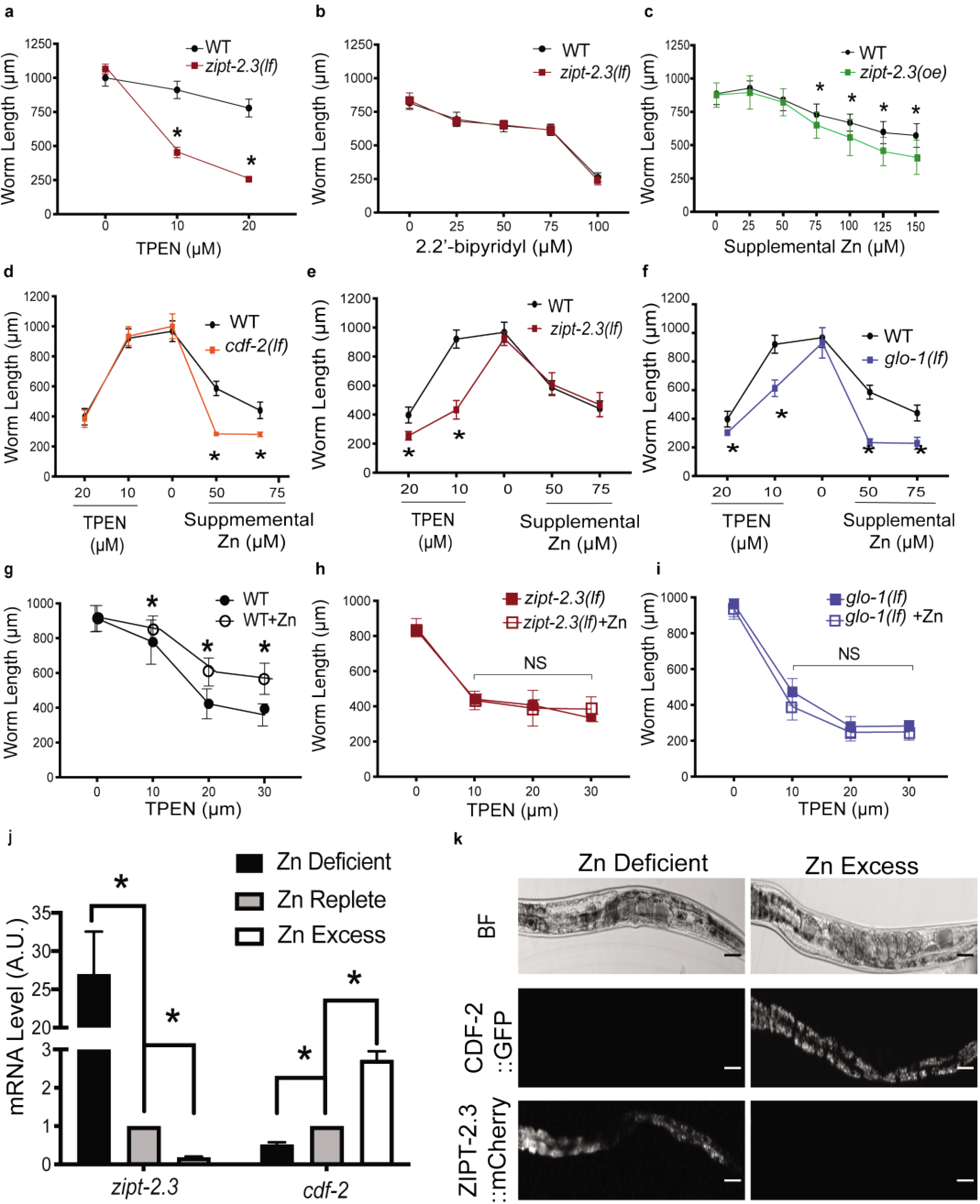
ZIPT-2.3 and CDF-2 function in zinc homeostasis and are regulated reciprocally by zinc levels. (a-f) L1 larvae were cultured on NAMM dishes containing the zinc chelator TPEN, the iron chelator 2,2’-bipyridyl, or supplemental zinc for three days, and the length of individual worms was measured. Values represent the average length +/- standard deviation (3 independent biological replicates, each with a minimum of twenty animals, *p<0.05). Genotypes: *zipt-2.3(ok2094)*, *amEx350* [*zipt-2.3(oe)*], *cdf-2(tm788), glo-1(zu391)*, and wild type. (g-i) L1 larvae were cultured on NAMM dishes containing 0 or 25µM (+Zn) supplemental zinc for 16 hours, shifted to NAMM dishes containing TPEN for three days, and analyzed for length. (j) A population of mixed-stage, wild-type animals were cultured with 200 µM supplemental zinc (zinc excess), 40 µM TPEN (zinc deficient), or 0µm supplemental zinc or TPEN (zinc replete) for 16 hours. RNA was analyzed by qPCR. The value in zinc replete conditions was set equal to 1.0 arbitrary units (AU), and other values normalized. Average of 3 biological replicates +/- standard deviation. (k) Transgenic L4 stage larvae expressing CDF-2::GFP and ZIPT-2.3::mCherry were cultured with 50 µM TPEN or 200 µM supplemental zinc for 16 hours. Representative images show one worm with bright field (BF, upper), green fluorescence (middle), or red fluorescence (lower). Scale bar = 10 μm.

The CDF-2 protein promotes storage in zinc excess conditions; *cdf-2(lf)* mutant animals displayed hypersensitivity to growth defects caused by high zinc toxicity but displayed normal growth in zinc deficient conditions (Fig. 2d). By contrast, *zipt-2.3(lf)* animals displayed hypersensitivity to growth defects caused by zinc deficiency but displayed normal growth in zinc excess conditions (Fig. 2e). To test the prediction that gut granules themselves are critical for zinc homeostasis, we examined animals with a mutation in *glo-1* that are defective in forming gut granules (21). *glo-1(lf)* animals were hypersensitive to growth defects caused by zinc deficiency and excess, demonstrating the central role of gut granules in zinc homeostasis (Fig. 2f).

When wild-type animals are exposed to excess zinc early in life, they display resistance to growth defects caused by zinc deficiency later in life, presumably because they mobilize stored zinc (Fig. 2g) (12). We predicted that *zipt-2.3* is necessary to take advantage of stored zinc. Consistent with this prediction, *zipt-2.3(lf)* mutants did not display increased resistance when exposed to excess zinc early in life (Fig. 2h). *glo-1(lf)* mutants displayed a similar defect, consistent with the central role of gut granules in zinc storage and release (Fig. 2i).

## Reciprocal regulation of ZIPT-2.3 and CDF-2 mediates zinc homeostasis

*cdf-2* mRNA levels increase in excess zinc conditions, and *zipt-2.3* mRNA levels increase in zinc deficient conditions (12, 19). To further analyze regulatory control, we cultured wild-type animals for 16 hours with 40µM TPEN (zinc deficient), 200µM supplemental zinc (zinc excess), or no supplemental zinc or TPEN (zinc replete) and analyzed mRNA. In zinc deficient conditions, the level of *zipt-2.3* mRNA was increased significantly and the level of *cdf-2* mRNA was decreased significantly compared to replete conditions. By contrast, in zinc excess conditions the level of *zipt-2.3* mRNA was decreased significantly and the level of *cdf-2* mRNA was increased significantly compared to replete conditions (Fig. 2j). Reciprocal regulatory control of CDF-2 and ZIPT-2.3 was also observed at the level of protein expression (Fig. 2k). These results suggest a mechanism for directional flow of zinc from the cytosol into the lumen of gut granules during excess, when CDF-2 levels are high and ZIPT-2.3 levels are low, and from the lumen of gut granules back into the cytosol during deficiency, when ZIPT-2.3 levels are high and CDF-2 levels are low.

## Super resolution microscopy reveals that gut granules are composed of two compartments that are remodeled in response to zinc excess and deficiency

If zinc homeostasis is regulated by shifting the ratio of CDF-2 and ZIPT-2.3, then mechanisms must exist to achieve dynamic changes in the composition of zinc transporters on the membranes of lysosome-related organelles. To define these mechanisms, we observed individual organelles at 120 nm resolution using super resolution microscopy. Animals raised in zinc replete medium were transferred to zinc deficient, replete, or excess conditions for 16 hours and visualized with three different fluorescent markers. One strain expressed CDF-2::GFP (labeled as red) and ZIPT-2.3::mCherry (labeled as green) and was stained with LysoTracker (labeled as blue) (Fig. 3a-d) (22). The second strain expressed CDF-2::mCherry (labeled as red) and was stained with LysoTracker (labeled as blue) and FluoZin-3 AM (labeled as yellow), which stains labile zinc (Fig. 3e-h). Individual organelles that did not overlap neighboring organelles were reconstructed in three dimensions, and maximum intensity projections (MIP) and line scans were used to estimate volumes and determine spatial relationships between markers. The MIP is a composite of all planes of a z-stack obtained during imaging, and thus represents the entire granule. For clarity of presentation, we applied a distinct arbitrary color for each marker: CDF-2 is red, ZIPT-2.3 is green, LysoTracker is blue, and labile zinc is yellow (Fig. 3-4, S3-4, 6-11).

**Figure 3.**
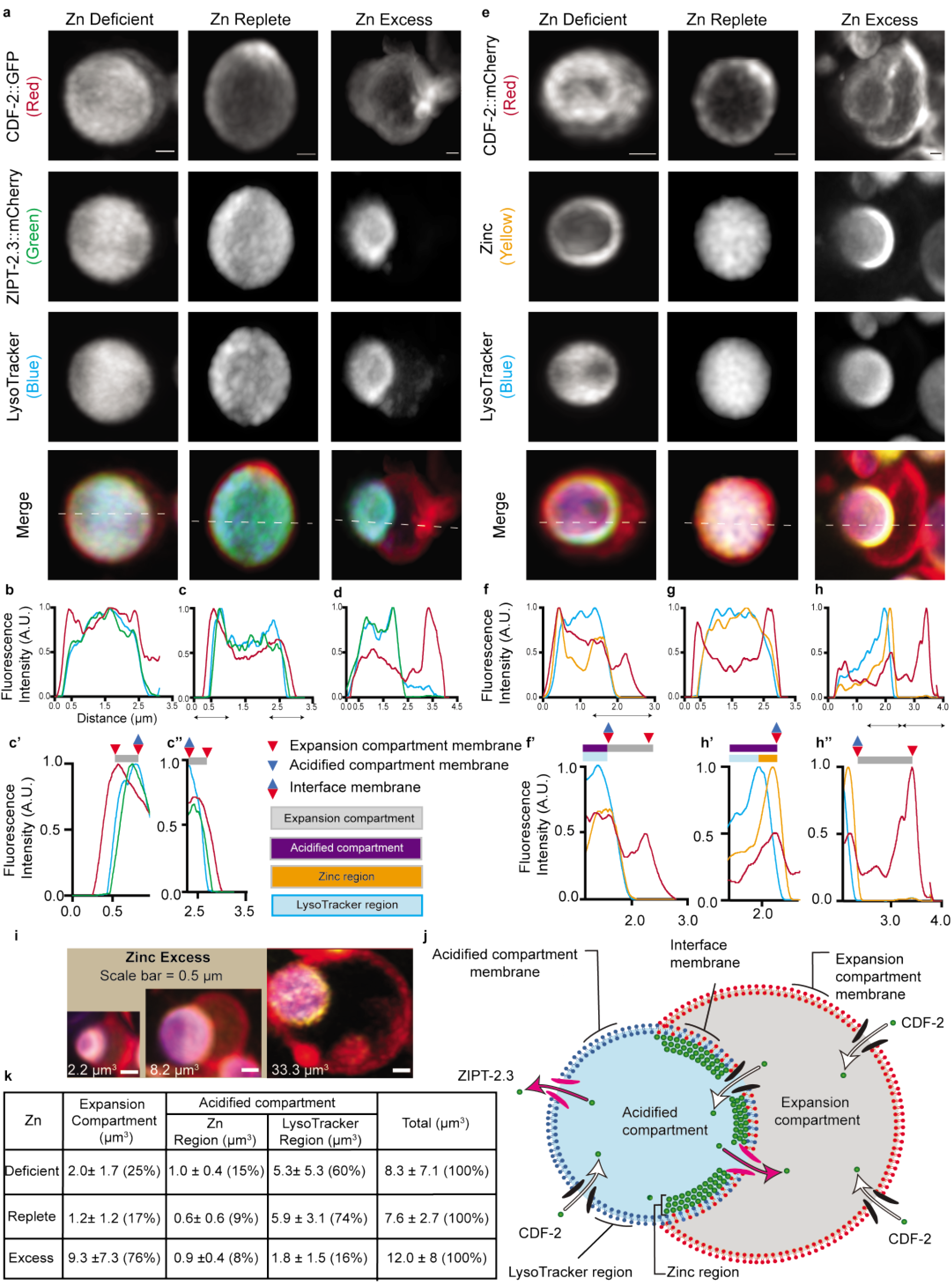
Super resolution microscopy reveals that gut granules have an acidified and an expansion compartment. (a) Transgenic L4 stage animals expressing CDF-2::GFP (true color green – arbitrary color red) and ZIPT-2.3::mCherry (true color red – arbitrary color green) were cultured for 16 hours in LysoTracker Blue (true color blue – arbitrary color blue) in either standard medium (Zn replete), 50 µM TPEN (Zn deficient) or 200 µM supplemental zinc (Zn excess). Individual gut granules were imaged by super resolution microscopy for green, red, and blue fluorescence, and a maximum intensity projection is displayed. Scale bar = 0.5 μm. (b-d) A line scan was performed, indicated by the dashed white line on merge image. For each color, the highest value was set equal to 1.0 arbitrary units (AU), and other values were normalized. (c’) Enlargements of specific regions indicated by black double-sided arrows. Annotations above indicate positions of membranes (triangles), compartments (purple and gray rectangles) and regions (blue and orange rectangles). (e-h) Transgenic L4 stage animals expressing CDF-2::mCherry (true color red – arbitrary color red) were cultured for 16-20 hours in LysoTracker blue (true color blue – arbitrary color blue) and the zinc dye FluoZin-3 AM (true color green – arbitrary color yellow). Culture conditions, imaging, and line scan analysis were similar to panel a-d. (i) Merge images of gut granules from animals cultured in zinc excess as in panel e illustrate volume variation (white number). Scale bar = 0.5 μm (j) Model of a gut granule in zinc excess conditions. Compartments, regions, and membranes are labeled. CDF-2 and ZIPT-2.3 proteins are black/white or pink arrows, respectively. (k) Volumes of the expansion compartment, zinc region, and LysoTracker region were calculated for gut granules from animals analyzed as in panel e. Values are average +/- S.D., and percent is the fraction of the total volume. N=11 deficient, 12 replete, and 11 excess.

**Figure 4.**
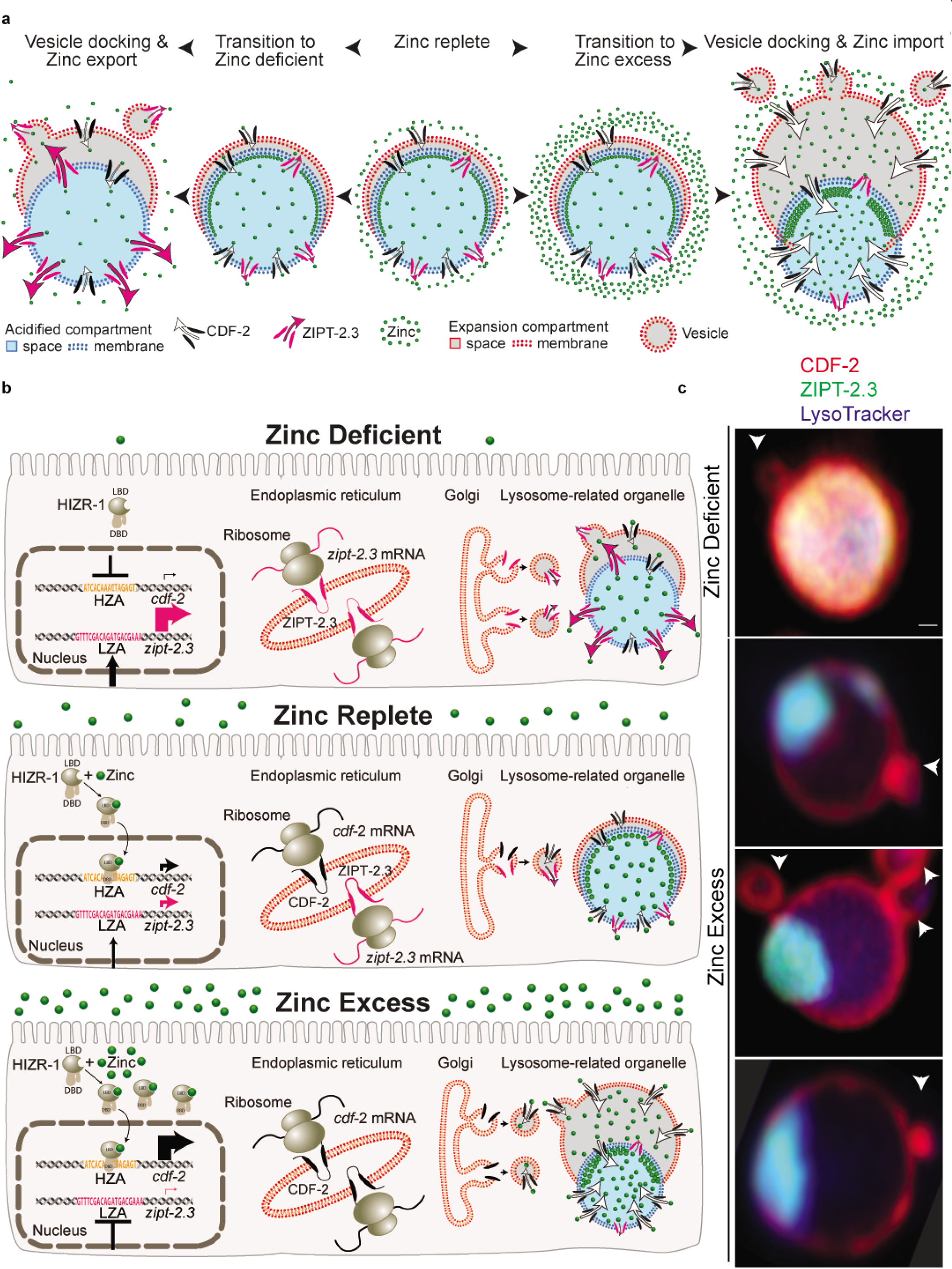
Vesicle fusion enlarges the expansion compartment and delivers CDF-2 and ZIPT-2.3 in zinc excess and deficient conditions, respectively. (a) Model of the morphological transition of gut granules and changes in transporter levels during the shift from zinc replete to zinc excess or deficient conditions. (b) Model of zinc homeostasis in zinc deficient, replete, and excess conditions. HIZR-1 and the HZA enhancer regulate expression of *cdf-2*; an undefined system for sensing low zinc and the LZA enhancer regulate expression of *zipt-2.3*. mRNA is translated in the ER, and Golgi-derived vesicles deliver CDF-2 and ZIPT-2.3 protein to gut granules, enlarging the expansion compartment in zinc excess and deficient conditions. (c) Transgenic L4 stage animals expressing CDF-2::GFP (true color green – arbitrary color red) and ZIPT-2.3::mCherry (true color red – arbitrary color green) were cultured for 16 hours in LysoTracker Blue (true color blue – arbitrary color blue) in either 50 µM TPEN (Zn deficient) or 200 µM supplemental zinc (Zn excess). Individual gut granules were imaged by super resolution microscopy for green, red, and blue fluorescence – these images are a three-color merge. Scale bar = 0.5 μm. White arrows indicate vesicles that contain CDF-2 and appear to be fusing with the expansion compartment membrane.

As depicted in Figure 3j, the results revealed that lysosome-related organelles are composed of two compartments in all zinc conditions: an **acidified compartment** that stains with LysoTracker, and an **expansion compartment** that is LysoTracker negative. The **acidified compartment membrane** contains both CDF-2 and ZIPT-2.3 and surrounds the spherical acidified compartment. The acidified compartment has two regions - the **LysoTracker region** forms the center of the sphere and stains strongly with LysoTracker and weakly with FluoZin-3 AM, whereas the **zinc region** forms the periphery of the sphere and stains strongly with FluoZin-3 AM but is depleted for LysoTracker staining. The zinc region appears as a crescent adjacent to the interface membrane in zinc excess conditions. The expansion compartment is a hemisphere that is dynamic in shape and volume. The **expansion compartment membrane** contains CDF-2 but not ZIPT-2.3 and appears to be attached to the acidified compartment membrane. The expansion compartment and the acidified compartment are separated by a portion of the acidified compartment membrane that we named the **interface membrane**.

To understand remodeling of compartments in response to changes in zinc levels, we measured compartment volumes. Volume calculations were based on the assumption that the compartments were spheres, hollow spheres, or hemispherical segments, which is based on our microscope images (Fig. S5). Our results below are summarized in Figure 4a.

In zinc replete conditions, gut granules are approximately spherical with a total average volume of ∼7.6 μm^3^ (Fig. 3e,k, S12a,b,i). The prominent acidified compartment represents 83% of the total volume, with a large LysoTracker region (∼5.9 μm^3^) and a small zinc region (∼0.6 μm^3^, Fig. S12e-h). The expansion compartment is contracted, with a total volume of ∼1.2 μm^3^, representing 17% of the total volume (Fig. S12,a-d). Although contracted, the expansion compartment can be visualized with super resolution microscopy in zinc replete conditions (Fig. 3a,S7). In many cases, the line scan reveals that the CDF-2 boundary (red line) is outside the ZIPT-2.3 boundary (green line), indicating the line scan passes through the expansion compartment membrane before passing through the acidified compartment membrane (Fig. 3c,c’,c’’). The ZIPT-2.3 membrane coincides closely with LysoTracker (Fig. S7). The LysoTracker and FluoZin-3 staining overlap extensively, with only a small zinc region outside the LysoTracker boundary (Fig. 3e). Overall, lysosome-related organelles in zinc replete medium appear to have stable membrane dynamics, store a small amount of zinc, and be primarily engaged in breaking down macromolecules in the prominent acidified compartment.

After 16 hours in zinc excess conditions, gut granules increase in total volume about 60% to an average of ∼12.0 μm^3^ (Fig. 3e,k). The prominent expansion compartment is shaped like a hemisphere and increases about 8-fold to ∼9.3 μm^3^; this represents 76% of the total volume (S12a-d). The acidified compartment is spherical and shrinks overall to 2.7 μm^3^, which is 24% of the total volume; this represents a large decrease in the LysoTracker region to 1.8 μm^3^ while the zinc region increases to ∼0.9 μm^3^. In most cases, the line scans reveal that the CDF-2 boundary (red line) is coincident with the ZIPT-2.3 boundary (green line) as it passes through the acidified compartment membrane (Fig. 3a,d, S3). In most cases there is a distinct zinc region shaped like a crescent: line scans reveal FluoZin-3 staining extends beyond the LysoTracker stain and is coincident with the CDF-2 membrane, indicating the crescent of zinc is in the acidified compartment rather than the expansion compartment (Fig. 3e,h’). Both the Lysotracker region and the zinc region are contained within the acidified compartment membrane and the interface membrane (Fig. 3,e,h-h’’, S11). Overall, lysosome-related organelles in zinc excess appear to have active membrane dynamics driving a growing expansion compartment and a large amount of zinc localized in a crescent shape; the structure suggests they are primarily engaged in zinc storage with a relatively small acidified compartment breaking down macromolecules.

After 16 hours in zinc deficient conditions, gut granules increase slightly to a total average volume of ∼8.3 μm^3^ (Fig. 3e,k). An expansion compartment shaped like a hemisphere is frequently visible with an average volume of ∼2.0 μm^3^; this is about 60% larger than in replete conditions, and it represents 25% of the total volume of the organelle. The acidified compartment is spherical and shrinks slightly to ∼6.3 μm^3^, which is 75% of the total volume; this represents a small decrease in the LysoTracker region to ∼5.3 μm^3^ while the zinc region slightly increases to ∼1.0 μm^3^ (Figure 3b,f,k). Overall, lysosome-related organelles in zinc deficiency appear to have active membrane dynamics leading to a larger expansion compartment and a mostly unchanged acidified compartment; the structure suggests they are primarily engaged in zinc release with an acidified compartment breaking down macromolecules.

In excess zinc conditions, the volume of gut granules varied ∼10-fold; of eleven analyzed in detail, the smallest was ∼2.2 μm^3^ and the largest was ∼33.3 μm^3^ (Fig. 3i, S11). Interestingly, the overall shape and proportions appeared to be similar despite these size differences. To rigorously determine how the proportions of gut granules scale with size, we analyzed the correlations between the volumes of the LysoTracker region, zinc region, and expansion compartment and the total volume. In zinc excess and deficiency, the expansion compartment and LysoTracker region positively correlated with the total volume, indicating that the gut granules have a similar composition regardless of size (Fig. S13). In zinc replete conditions, only the LysoTracker region positively correlated with total volume (Fig. S13).

## Vesicles appear to deliver zinc transporters to lysosome-related organelles and mediate the volume change of the expansion compartment

In zinc excess conditions, we frequently observed small, spherical vesicles that were positive for CDF-2 adjacent to or fusing with the expansion compartment. In zinc deficient conditions, we occasionally observed such vesicles (Fig. 4c). Based on these observations we propose that vesicle fusion is responsible for delivering CDF-2 to gut granules in zinc excess conditions and is the source of the increase in the extent of the expansion compartment membrane. Similarly, we propose that vesicle fusion is responsible for delivering ZIPT-2.3 to gut granules in zinc deficient conditions and is the source of the increase in the extent of the expansion compartment membrane. The process seems to be robust in zinc excess conditions, resulting in the appearance of many vesicles and a dramatic increase in the volume of the expansion compartment; the process is less robust in zinc deficient conditions, since fewer vesicles were observed and the change in the expansion compartment is subtler (Fig. 4a).

Based on these observations and previous studies of zinc-regulated transcription, we propose an integrated model of zinc homeostasis (Fig. 4b). In zinc excess conditions, high levels of cytoplasmic zinc lead to activation of the high zinc sensor HIZR-1(23). When zinc binds the HIZR-1 ligand-binding domain (LBD), HIZR-1 translocates to the nucleus where its DNA binding domain (DBD) interacts with the High Zinc Activation (HZA) enhancer, increasing *cdf-2* transcription. By contrast, transcription of *zipt-2.3* is decreased by a mechanism that has not been established. Increased levels of *cdf-2* transcripts result in increased translation of CDF-2 protein in the endoplasmic reticulum and the generation of vesicles that fuse with the expansion compartment of gut granules. Vesicle fusion adds membrane and increases the volume of the expansion compartment, and the increased levels of CDF-2 promote zinc transport and detoxification. Zinc that is imported into the expansion compartment or the acidified compartment is concentrated in the zinc region (Fig. 4b, lower). In zinc replete conditions, transcription of *cdf-2* and *zipt-2.3* are balanced, and only a small number of vesicles fuse with gut granules, so the expansion compartment is contracted (Fig. 4b middle). In zinc deficient conditions, the Low Zinc Activation (LZA) enhancer is activated, leading to increased levels of *zipt-2.3* transcripts. By contrast the *cdf-2* promoter is repressed by an unknown mechanism. Increased levels of *zipt-2.3* transcripts result in increased translation of ZIPT-2.3 protein in the endoplasmic reticulum and the generation of vesicles that fuse with the expansion compartment of gut granules. Vesicle fusion enlarges the expansion compartment slightly, and the increased levels of ZIPT-2.3 promote zinc export (Fig. 4b upper).

## Discussion

CDF-2 was previously identified as the transporter that stores zinc in gut granules (12), but the mechanism of release was not defined. Here we identify ZIPT-2.3 as the zinc transporter that mediates release of stored zinc from gut granules. Cell-based assays demonstrated that ZIPT-2.3 protein transports zinc, microscopy studies showed specific localization in lysosome-related organelles, and analysis of loss-of-function and gain-of-function mutants documented multiple phenotypes consistent with a role in mobilizing stored zinc. This is an important advance because it defines the pair of transporters that mediate zinc storage and release.

ZIPT-2.3 and CDF-2 display dramatic and reciprocal regulation in response to zinc conditions, identifying one mechanism for directional storage and release. In the transition from zinc deficient to zinc excess conditions, *cdf-2* transcript levels increase ∼6 fold and *zipt-2.3* transcript levels decrease ∼130 fold. Increased transcription of *cdf-2* in zinc excess conditions is mediated by the HIZR-1 nuclear receptor transcription factor and the HZA enhancer, and increased transcription of *zipt-2.3* in zinc deficient conditions is mediated by the LZA enhancer. The mechanism of *cdf-2* repression in zinc deficiency and *zipt-2.3* repression in zinc excess have not been defined, and the identification of these regulatory events establishes the foundation for future studies to define these control mechanisms.

Lysosome-related organelles are typically considered to be spherical and surrounded by a single lipid bilayer. Roh *et al.* (2012) first identified bilobed granules in zinc excess using the CDF-2 marker and confocal microscopy. However, these studies did not characterize the structure in detail or establish its function. Here we used super resolution microscopy to gain important new insights into this structural feature of lysosome-related organelles. Whereas Roh *et al.* (2012) suggested bilobed granules were a specialization for high zinc, here we demonstrate that the expansion compartment is a permanent structural feature of lysosome-related organelles. In zinc replete conditions, the expansion compartment is contracted and difficult to appreciate with standard confocal microscopy but detectable by super-resolution techniques. Furthermore, the expansion compartment inflates in both zinc excess and deficiency, indicating it is not a specific adaptation for one zinc extreme but rather a response to changes in zinc levels. Whereas Roh *et al.* (2012) suggested that labile zinc accumulates in the expansion compartment during zinc excess, super-resolution microscopy reveals that labile zinc accumulates in the acidified compartment, not the expansion compartment, and it is concentrated in a crescent at the interface membrane. The mechanism that directs stored zinc to this location is unknown. Roh *et al.* (2012) did not establish how the expansion compartment increases in volume. The results presented here suggest the expansion compartment increases in volume by fusion of vesicles containing CDF-2 in zinc excess, thereby increasing the capacity for zinc storage and detoxification, and by the fusion of vesicles containing ZIPT-2.3 in zinc deficiency, thereby increasing the capacity for zinc mobilization, Whereas Roh *et al.* (2012) did not establish the function of the expansion compartment, the results presented here indicate that the expansion compartment allows rapid changes in the composition of zinc transporters while preserving the pH of the acidified compartment by providing a separate compartment for vesicle fusion. The enhanced resolution of these microscopy techniques combined with the identification of CDF-2 and ZIPT-2.3 provide surprising new insights into the structure of lysosome-related organelles and mechanisms of zinc homeostasis, and they raise a new set of fascinating questions. Are all transmembrane proteins delivered to the expansion compartment membrane, or are some delivered directly to the acidified compartment membrane? What mechanism allows CDF-2 to localize to the membrane of both compartments, whereas ZIPT-2.3 localizes specifically to the acidified compartment membrane? How is zinc concentrated next to the interface membrane? Is the expansion compartment an evolutionarily conserved structural feature of lysosome-related organelles in other species that store zinc in these structures? In humans, ZIP8 releases zinc stored in lysosomes of T cells (23), and ZIP13, a gene implicated in the connective tissue disorder spondylocheiro dysplastic Ehlers-Danlos syndrome, releases zinc from intracellular vesicles (24, 25). Thus, ZIP proteins likely play a conserved role in releasing zinc, and the new paradigm for zinc regulation through organelle remodeling described here might also be conserved. These results highlight that lysosome-related organelles are multifunctional – in addition to the canonical function of macromolecule degradation in the acidified compartment, they function as a site of zinc storage, and the expansion compartment is an unexpected structural feature that promotes this dual function.

## Acknowledgments

We thank Laura Kyro for graphics and Suzanne Pfeffer for advice.

## Author Contributions

Conceptualization: A.D.M, N.D., and K.K.; Methodology: A.D.M., N.D., J.T., and K.K.; Formal Analysis: A.D.M., N.D., J.T., and K.K.; Investigation: A.D.M., N.D., J.T., C.C., D.H., J.K., Z.P., D.L.S.; Writing-Original Draft: A.D.M., N.D., and K.K., Writing-Review & Editing: A.D.M., N.D., J.T., C.C., D.H., J.K., Z.P., D.L.S., and K.K.; Visualization: A.D.M., N.D., and J.T.; Supervision: K.K.; Funding acquisition: K.K..

## Data and materials availability

All data is available in the manuscript or supplementary materials.

## Experimental Procedures

### General Methods and Strains

*C. elegans* strains were cultured at 20°C on nematode growth medium (NGM) dishes with a lawn of *E. coli* OP50 unless otherwise noted (28). The Bristol N2 strain was wild type and parental strain of all mutants. The following mutations and transgenes were used: *zipt-2.3 (ok2094) II* (*29*), *cdf-2(tm788) X* (19), *glo-1(zu391) X* (*21*), *amEx132(cdf-2::mCherry;rol-6^D^)*(*12*)*, amIs4(cdf-2::GFP::unc-119(+)*. The following transgenic strains were generated for this study: WU1816 (*zipt-2.3p::ZIPT-2.3::mCherry(amEx348*)), WU1824 (*ges-1p::zipt-2.3::T7*(*amEx350*)), WU1984 *cdf-2(tm788);amIs4; zipt-2.3p::ZIPT-2.3mCherry (amEx191),* and *zipt-2.3(ok2094);zipt-2.3p:: ZIPT-2.3::mCherry (amEx348)*.

### Measuring worm growth with metal excess or chelation

Gravid adult hermaphrodites were treated with bleach and sodium hydroxide, eggs were incubated in M9 solution overnight to allow hatching and synchronized arrest at the L1 larval stage, and L1 animals were transferred to noble agar minimum media (NAMM) dishes (18). For metal deficiency studies, NAMM was supplemented with N,N,N’,N’-tetrakis(2-pyridylmethyl)ethane-1,2-diamine (TPEN, Sigma-Aldrich), a zinc-specific chelator, 2,2-bipyridyl (Sigma-Aldrich), an iron specific chelator, or 1,2-Diaminocyclohexanetetraacetic acid monohydrate (Sigma-Aldrich), a manganese specific chelator. For zinc excess studies, NAMM was supplemented with ZnSO_4_ (Sigma Aldrich). Dishes were seeded with 5x concentrated *E. coli*. After culturing for 3 days, animals were paralyzed in a 10 mM sodium azide solution in M9 and mounted on a 2% agarose pad on a microscope slide. Images were captured using a Zeiss Axioplan 2 microscope equipped with a Zeiss AxioCam MRm digital camera. Lengths of individual animals were measured using ImageJ software by drawing a line from the nose to the tip of the tail of each animal.

To analyze growth after a short period of exposure to excess zinc (Fig. 2G-I), we bleached gravid adults to obtain arrested L1 larvae as described above. These animals were cultured on NAMM dishes supplemented with either 0 or 25µM zinc for 16 hours. Animals were washed with M9 containing 0.01% Tween-20, cultured on NAMM dishes containing 0, 10, 20, or 30 µM TPEN seeded with 5x concentrated *E. coli* OP50 for 3 days, and the length of each animal was determined as described above.

### Spinning Disk Microscopy

Transgenic L4 stage animals expressing CDF-2::GFP and ZIPT-2.3::mCherry were cultured for 16 hours in LysoTracker blue on standard NAMM dishes or dishes containing 50 μM TPEN or 200 μM ZnSO_4_. LysoTracker Blue (Invitrogen) was diluted in *E. coli* OP50 to obtain a concentration of 1 μM, respectively, and dispensed on either zinc deficient, replete, or excess dishes. Animals were anesthetized in 50 μM NaN_3_, mounted on an agar pad, and sealed with a coverslip. Microscopy was performed with the Nikon Spinning Disk confocal microscope using the 405, 488, and 561 laser lines to detect, LysoTracker Blue, CDF-2::GFP, and ZIPT-2.3::mCherry, respectively. All images were captured using the 60x objective.

### Zinc Uptake Assay

Zinc uptake assays were performed as described in Zhao *et al.* (2018) with the same set of controls (14). Briefly, HEK293T cells were seeded on Poly-D-lysine coated 24-well plates (Corning). The next day the cells were transfected with a plasmid encoding ZIPT-2.3 or pcDNA-3.1(+) (a vector only control) using Lipofectamine 2000 (Invitrogen). After 48 hours, cells were washed once with pre-warmed uptake buffer (15 mM HEPES, 100 mM glucose, 150 mM KCl, pH 7.0) and incubated for 15 minutes in pre-warmed uptake buffer that contained the radioactive tracer 65 ZnCl_2_ (PerkinElmer) and non-radioactive ZnCl_2_ (Sigma). Uptake was halted by applying the same volume of ice-cold stop buffer (15 mM HEPES, 100 mM glucose, 150 mM KCl, 1 mM EDTA, pH 7.0). Cells were gently washed with ice-cold stop buffer twice and disassociated with trypsin. Radioactivity incorporated into the cells was measured with a Beckman LS 6000 Scintillation Counter. In parallel experiments conducted without adding metals, the cells were lysed with lysis buffer (2 mM Tris-HCl, 150 mM NaCl, 1% Triton X-100), and protein levels were measured with the Bio-Rad DC protein assay. 65 Zn uptake was normalized to total protein measured in this parallel assay. The data shown in Figure 1b are typical of multiple independent experiments.

### Plasmid DNA construction and transgenic strain generation

To generate an epitope tagged construct expressing *zipt-2.3,* we used *C. elegans* wild-type genomic DNA as a template, and the polymerase chain reaction (PCR) was used to amplify DNA fragments with Phusion polymerase (New England Biolabs) of the genomic sequence using a forward primer that starts from 2199 bases upstream of the ATG start codon of *zipt-2.3* and a reverse primer that contained the codon preceding the stop codon of *zipt-2.3* and the coding sequence of the T7 epitope (MASMTGGQQMG). Amplified DNA was ligated into pBluescript SK+ along with the DNA of the *unc-54* 3’ untranslated region. The mCherry plasmid (pND32) was generated by amplifying the promoter and coding region of *zipt-2.3* by PCR and cloning into the plasmid pSC6, which contains mCherry upstream of the *unc-54* 3’ UTR. To overexpress *zipt-2.3* we replaced the promoter of *zipt-2.3* by using PCR to amplify 2100 bases upstream of the ATG start codon of *ges-1* representing the promoter, which was ligated into the plasmid containing the coding region of *zipt-2.3*, the T7 epitope, and the *unc-54* 3’ UTR (30). All plasmid sequences were confirmed by standard DNA sequencing. To generate transgenic strains, we injected plasmids into N2 animals and selected animals that displayed the co-injection marker phenotype (31).

### Zinc shift assays with FluoZin-3

FluoZin-3 acetoxymethyl (AM) ester (excitation 494 nm, emission 516 nm) (Molecular Probes) was reconstituted in dimethylsulfoxide (DMSO) to generate a 1mM stock solution. This solution was diluted in 5X concentrated *E. coli* OP50 to generate a final concentration of 20µM, which was dispensed on NAMM dishes. L4 stage hermaphrodites were cultured on these dishes supplemented with 200µM zinc for 16 hours in the dark, and transferred to NGM dishes with no FluoZin-3 AM for 30 minutes to reduce the amount of dye within the intestinal lumen. These animals were examined for fluorescence by mounting into a 10mM sodium azide solution in M9 placed on a 2% agarose pad on a microscope slide. The animals were imaged with a Zeiss Axioplan 2 microscope equipped with a FITC filter, and a Zeiss AxioCam MRm digital camera using identical settings and exposure times. The intestine on the anterior part of each animal was analyzed. Animals were then transferred to NAMM dishes with FluoZin-3 AM and 0 or 100µM TPEN, and the fluorescence intensity (in arbitrary units) of the anterior intestines in each condition were measured using FIJI (32).

### Quantitative real-time PCR (qRT-PCR)

We performed qRT-PCR as previously described with minor modifications (19). To analyze transcript levels of all 14 *zipt* genes, WT animals were cultured in excess zinc (Fig. S2). To analyze transcript levels of zinc responsive genes in WT, *zipt-2.3(lf),* and *zipt-2.3(oe)* strains (Fig. 2), we collected mixed-stage populations of *C. elegans* by washing and cultured them for 16 hours on NAMM dishes seeded with concentrated *E. coli* OP50 and supplemented with 0µM or 40µM TPEN or 200µM zinc sulfate. Animals were collected by washing, and RNA was isolated using the TRIzol reagent (Invitrogen) and treated with DNase I. cDNAs were synthesized using the High Capacity cDNA Reverse Transcription kit according to the manufacturer’s protocol (Applied Biosystems). PCR was performed using an Applied Biosystems 7900 thermocycler and iTaq Universal SYBR Green Supermix (Bio-Rad). In all cases, the transcript level was normalized to the transcript level of a reference gene (*ama-1*) in the same sample. Fold change was determined by dividing the normalized transcript level at 200µM supplemental zinc or 40µM TPEN by the normalized transcript level at 0µM supplemental zinc or TPEN (Fig. S2).

### Super resolution Microscopy

Transgenic L4 stage animals expressing CDF-2::GFP and ZIPT-2.3::mCherry were cultured for 16 hours in LysoTracker blue on standard NAMM dishes or dishes containing 50 μM TPEN or 200 μM ZnSO_4_. Transgenic L4 stage animals expressing CDF-2::mCherry were cultured for 16-20 hours in LysoTracker Blue and the zinc dye FluoZin-3 AM. FluoZin-3 AM and LysoTracker Blue were diluted into *E. coli* OP50 to obtain a final concentration of 10 μM and 1 μM, respectively, and dispensed on either zinc deficient, replete, or excess dishes. Animals were anesthetized in 50 μM NaN_3_, mounted on an agar pad, and sealed with a coverslip. Superresolution microscopy was performed with the Zeiss LSM 880 Confocal with Airyscan. Gut granules were selected for analysis if they displayed fluorescence from LysoTracker Blue, FluoZin-3, and CDF-2::mCherry or if they displayed fluorescence from LysoTracker Blue, ZIPT-2.3::mCherry, and CDF-2::GFP. Images of gut granules were captured in z-stack using the 60x objective. FluoZin-3 and CDF-2::GFP were detected using the 488 nm laser, LysoTracker Blue was detected using the 405 nm laser, and ZIPT-2.3::mCherry and CDF-2::mCherry were detected using the 561 nm laser. Images were deconvolved using AiryScan processing to achieve 120 nm resolution.

### Image Analysis & Volume Calculations

Post-imaging analysis was performed with Imaris software (Bitplane) and FIJI. Individual gut granules were cropped and isolated. Arbitrary colors were used for display images as follows: CDF-2::GFP and CDF-2::mCherry (red), ZIPT-2.3::mCherry (green), LysoTracker Blue (blue), and FluoZin-3 (yellow).

#### Line scan

The length of the entire granule was traced across end to end and is indicated by a dashed line. The line captures the distribution of membranes and compartment spaces over the length of the granule. Line colors corresponded to the arbitrary colors above.

#### Granule volumes

The diameter of each compartment was measured by tracing the distance across each compartment end to end three times and calculating the average value. The volume of the LysoTracker region was calculated based on the assumption that it is spherical, which appears to be true in all zinc conditions. We measured the diameter, calculated the radius as 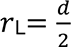, and calculated the volume as 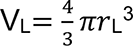 Figure S5a-c). The zinc region surrounds the LysoTracker region, and the volume was calculated based on the assumption that it is a hollow sphere. We measured the diameter, calculated the radius as 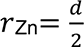, and calculated the volume as 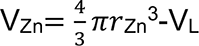 (Figure S5a’-c’). The expansion compartment has a more complex morphology that depends on zinc conditions. In zinc replete condition, the expansion compartment surrounds the zinc region and is not inflated, and the volume was calculated based on the assumption that it is a hollow sphere. We measured the diameter, calculated the radius as 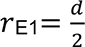, and calculated the volume as 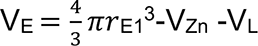 (Figure S5b’’). In zinc deficient and excess conditions, the expansion compartment is inflated, and we calculated the volume by separately determining (1) the volume of the region surrounding the acidified compartment as described above and (2) the volume of the inflated region. In zinc excess conditions, we assumed the inflated region is a sphere, measured the diameter, calculated the radius as 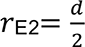, and calculated the volume as 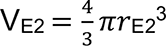. The total volume of the expansion compartment is 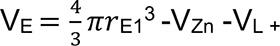 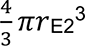 (Figure S5c’’). In zinc deficient conditions, we assumed the inflated region is a hemisphere, measured the radius as 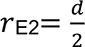, measured the height as h_E2_, and calculated the volume of the hemisphere using the formula 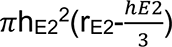 The total volume of the expansion compartment is 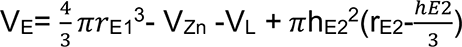 (Figure S5a’’).

## Statistical analysis

Comparisons of data were performed using the two-tailed unpaired Student’s t-test, and a P value <0.05 was considered statistically significant. Correlation analysis was conducted by Pearson correlation, and a P value <0.05 was considered statistically significant. R values >0 indicate a positive correlation, R <0 indicate a negative correlation, and R=0 indicates no correlation. Correlations were first analyzed with a 1-way ANOVA; when P<0.05, data were further analyzed by a student’s t-test.

---

**Supplemental Figure 1.**
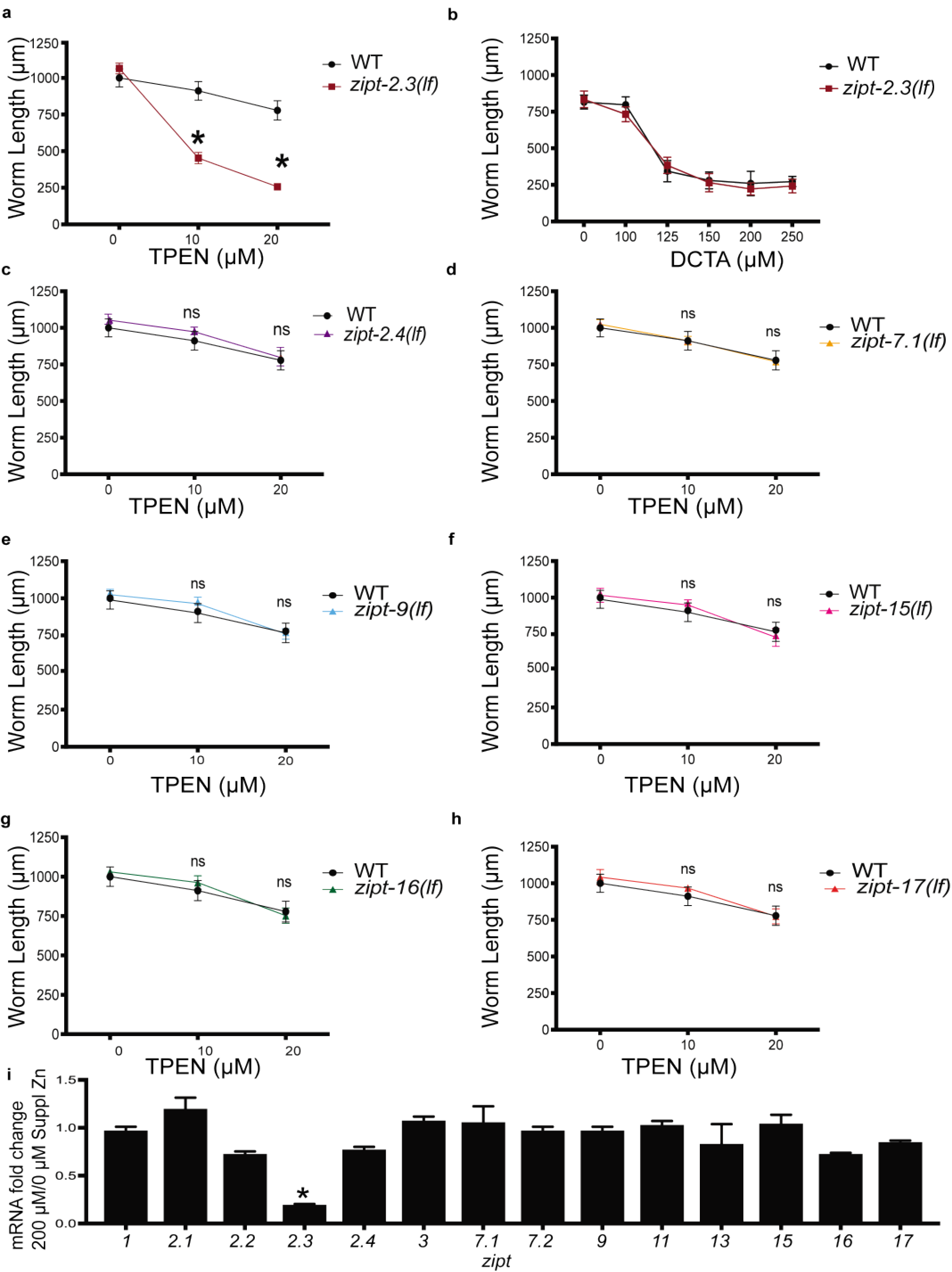
(with main Fig. 2). *zipt-2.3(lf)* hypersensitivity to zinc deficient conditions, and *zipt-2.3* transcriptional repression in zinc excess conditions were not displayed by other *zipt* genes. (a-h) Animals were synchronized at the L1 larval stage and cultured for three days on standard NAMM dishes or dishes supplemented with the zinc chelator TPEN or the manganese chelator DCTA. Length of individual worms was measured using microscopy to capture images and ImageJ software for analysis. Values represent the average length +/- standard deviation (3 independent biological replicates, each with a minimum of twenty animals, *P<0.05). Genotypes were wild type, *zipt-2.3(ok2094) II, zipt-2.4(ok2221) I, zipt-7.1(ok971) IV, zipt-9(ok876) V, zipt-15(ok2160) IV, zipt-16(ok875) V,* and *zipt-17*(*ok745) IV*. Panel a is identical to Figure 2a and is shown here to facilitate comparisons. (i) Wild-type animals were cultured with 0 or 200 µM supplemental zinc for 16 hours, and mRNA levels were analyzed by qPCR for all 14 *C. elegans zipt* genes. Values are the ratio of mRNA levels at 200µM supplemental zinc/0µM supplemental zinc, an indication of transcriptional regulation by high zinc. Average of 3 biological replicates +/- standard deviation (*P<0.05). Only *zipt-2.3* mRNA levels were significantly reduced in excess zinc, indicating this is a specific regulatory response.

**Supplemental Figure 2.**
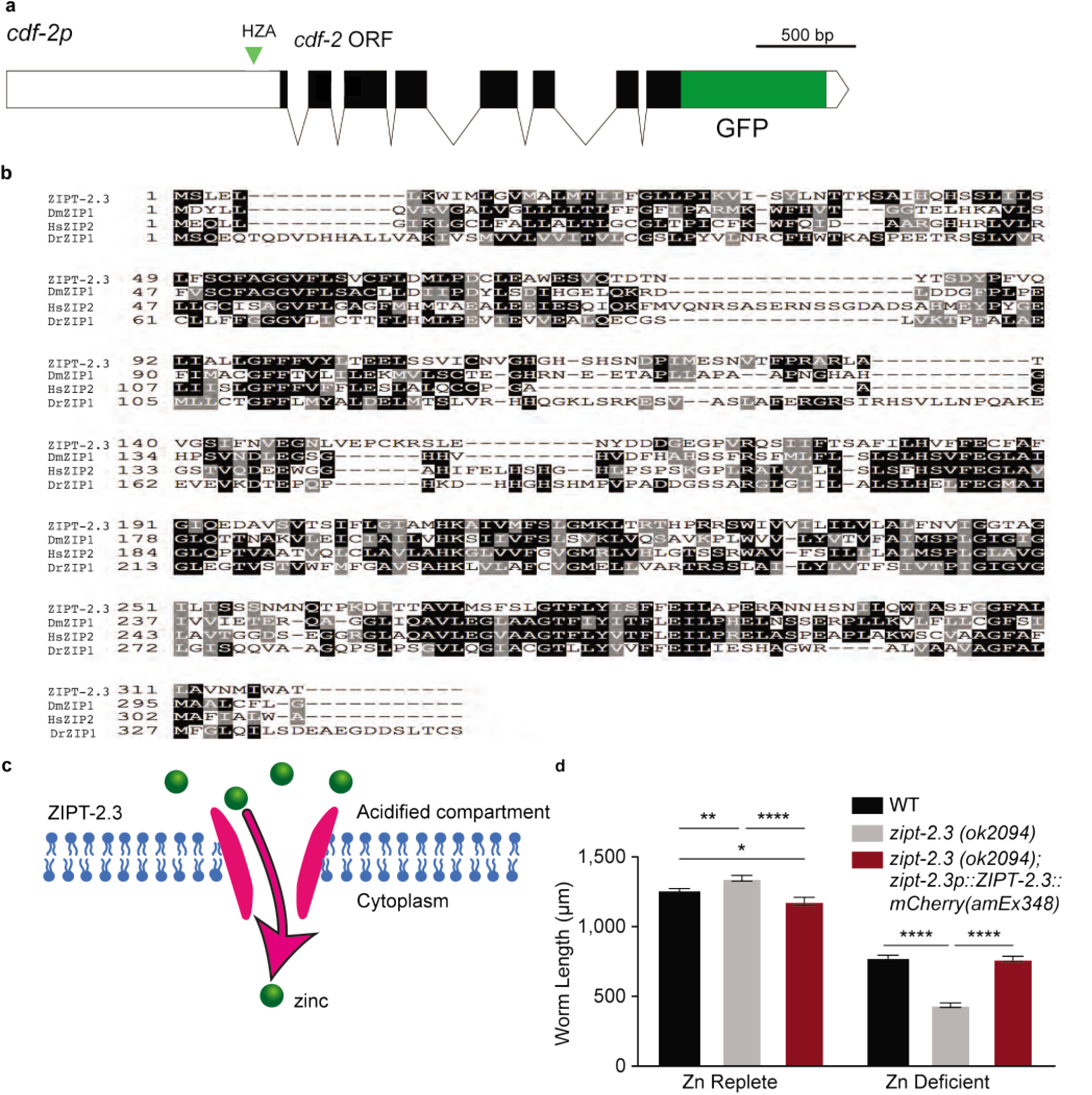
(with main Fig. 1). ZIPT-2.3 is a conserved zinc transporter. (a) The diagram shows a portion of a plasmid in transgenic strains that express CDF-2::GFP (*amIs4).* White boxes represent promoter and 3’ UTR regions, black boxes and lines represent CDF-2 coding regions and introns, green represents the GFP coding region, and the green triangle represents the HZA enhancer. (b) An alignment of the predicted *C. elegans* ZIPT-2.3, *D. melanogaster* ZIP1, human ZIP2, and *D. rerio* ZIP1 proteins. Identical and similar amino acids are highlighted in black and grey, respectively. (c) Model of the ZIPT-2.3 transporter (pink bars), a predicted transmembrane protein, in a lipid bilayer (blue). ZIPT-2.3 transports zinc (green circles) from the acidified compartment of gut granules to the cytoplasm (pink arrow). (d) Wild-type, *zipt-2.3(ok2094)*, and *zipt-2.3(ok2094);zipt-2.3p:: ZIPT-2.3::mCherry (amEx348)* animals were synchronized at the L1 larval stage and cultured for three days on standard NAMM dishes (zinc replete) or dishes supplemented with 50 μM TPEN (zinc deficient). Length of individual worms was measured using microscopy to capture images and ImageJ software for analysis. Values represent the average length +/- standard error (3 independent biological replicates, and each replicate included a minimum of twenty animals, *p<0.05, **p<0.01, ***p<0.001). *zipt-2.3(ok2094)* animals displayed significantly reduced growth in zinc deficient conditions, and this defect was rescued by expression of the ZIPT-2.3::mCherry protein, indicating this fusion protein is functional *in vivo*.

**Supplemental Figure 3.**
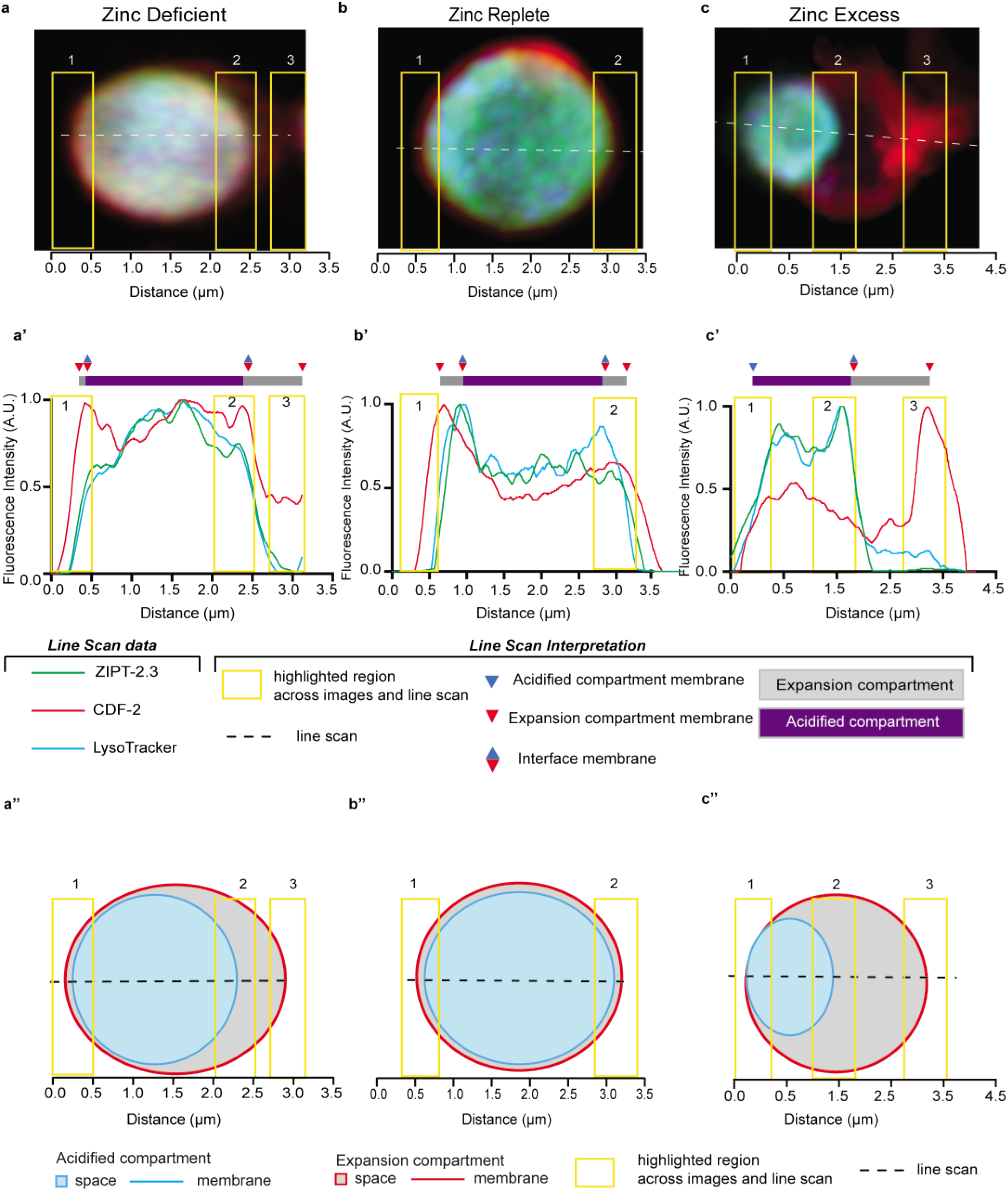
(with main Fig. 3a-d). Analysis of super resolution microscopy images using line scans. Transgenic L4 stage animals expressing CDF-2::GFP (true color green – arbitrary color red) and ZIPT-2.3::mCherry (true color red – arbitrary color green) were cultured for 16 hours in LysoTracker Blue (true color blue – arbitrary color blue) in either standard medium (Zn replete), 50 µM TPEN (Zn deficient) or 200 µM supplemental zinc (Zn excess). Individual gut granules were imaged by super resolution microscopy for green, red, and blue fluorescence. (a-c) Images display a three-color merge of a maximum intensity projection. A line scan was performed (dashed white line on merge image). Yellow boxes labeled 1-3 indicate sections of the line scan that cross membranes. (a’-c’) For each color in the line scan, the highest value was set equal to 1.0 arbitrary units (AU), and other values were normalized. Annotations above indicate positions of membranes (triangles) and compartments (purple and gray rectangles). Yellow boxes correspond to the images above and cartoons below. The same images and line scans are shown in Figure 3a-d. (a’’-c’’) Cartoons of lysosome-related organelles. The expansion compartment space is gray and the membrane is a red line. The acidified compartment space is blue and the membrane is a blue line. The dashed black line indicates the line scan, and yellow boxes correspond to the images above.

**Supplemental Figure 4.**
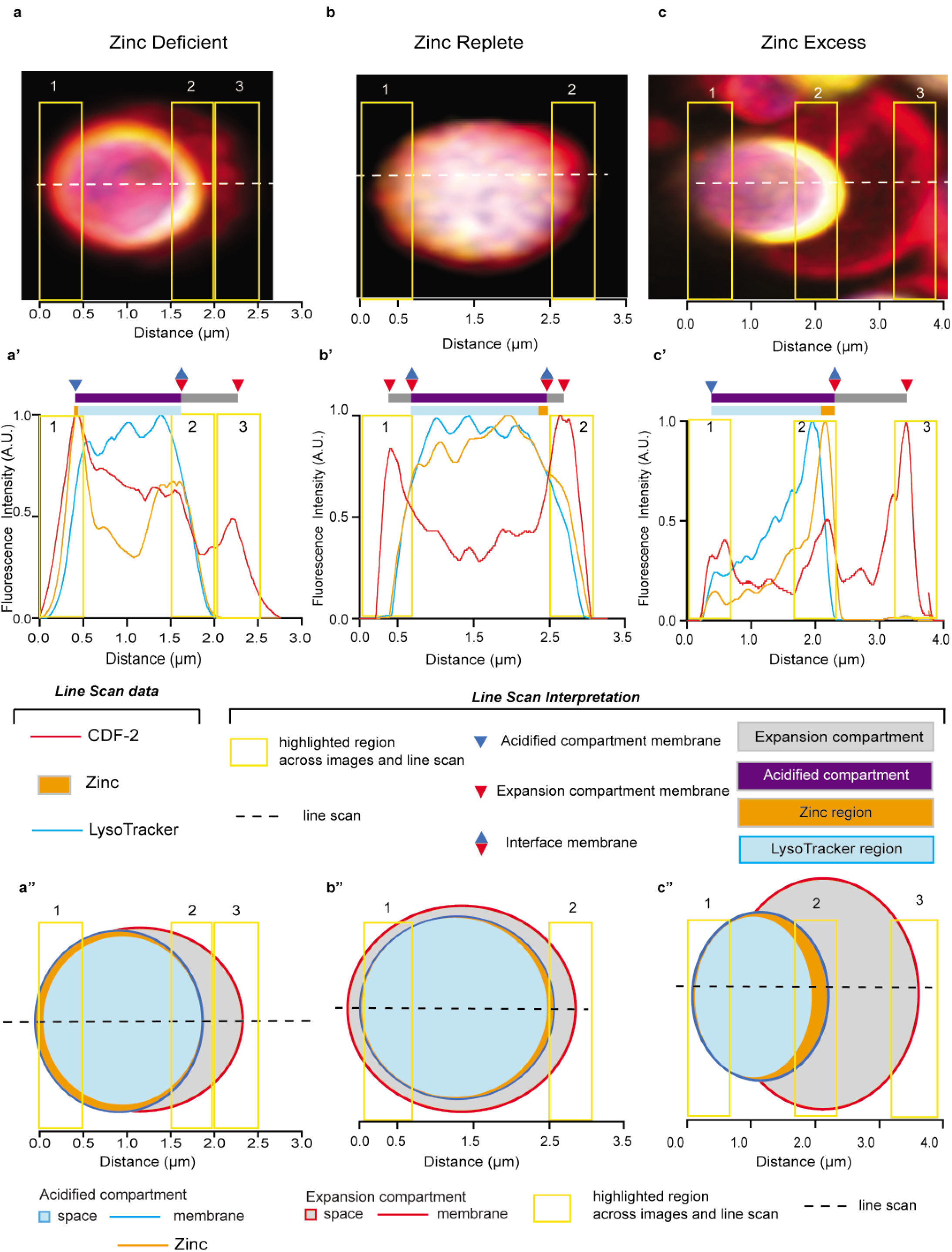
(with main Fig. 3e-h). Analysis of super resolution microscopy images using line scans. Transgenic L4 stage animals expressing CDF-2::mCherry (true color red – arbitrary color red) were cultured for 16-20 hours in LysoTracker blue (true color blue – arbitrary color blue) and the zinc dye FluoZin-3 AM (true color green – arbitrary color yellow) in either standard medium (Zn replete), 50 µM TPEN (Zn deficient) or 200 µM supplemental zinc (Zn excess). Individual gut granules were imaged by super resolution microscopy for green, red, and blue fluorescence. (a-c) Images display a three-color merge of a maximum intensity projection. A line scan was performed (dashed white line on merge image). Yellow boxes labeled 1-3 indicate sections of the line scan that cross membranes. (a’-c’) For each color in the line scan, the highest value was set equal to 1.0 arbitrary units (AU), and other values were normalized. Annotations above indicate positions of membranes (triangles), compartments (purple and gray rectangles), and regions (blue and orange rectangles). Yellow boxes correspond to the images above and cartoons below. The same images and line scans are shown in Figure 3e-h. (a’’-c’’) Cartoons of lysosome-related organelles. The expansion compartment space is gray, and the membrane is a red line. The acidified compartment space is blue for the LysoTracker region and orange for the zinc region, and the membrane is a blue line. The dashed black line indicates the line scan, and yellow boxes correspond to the images above.

**Supplemental Figure 5.**
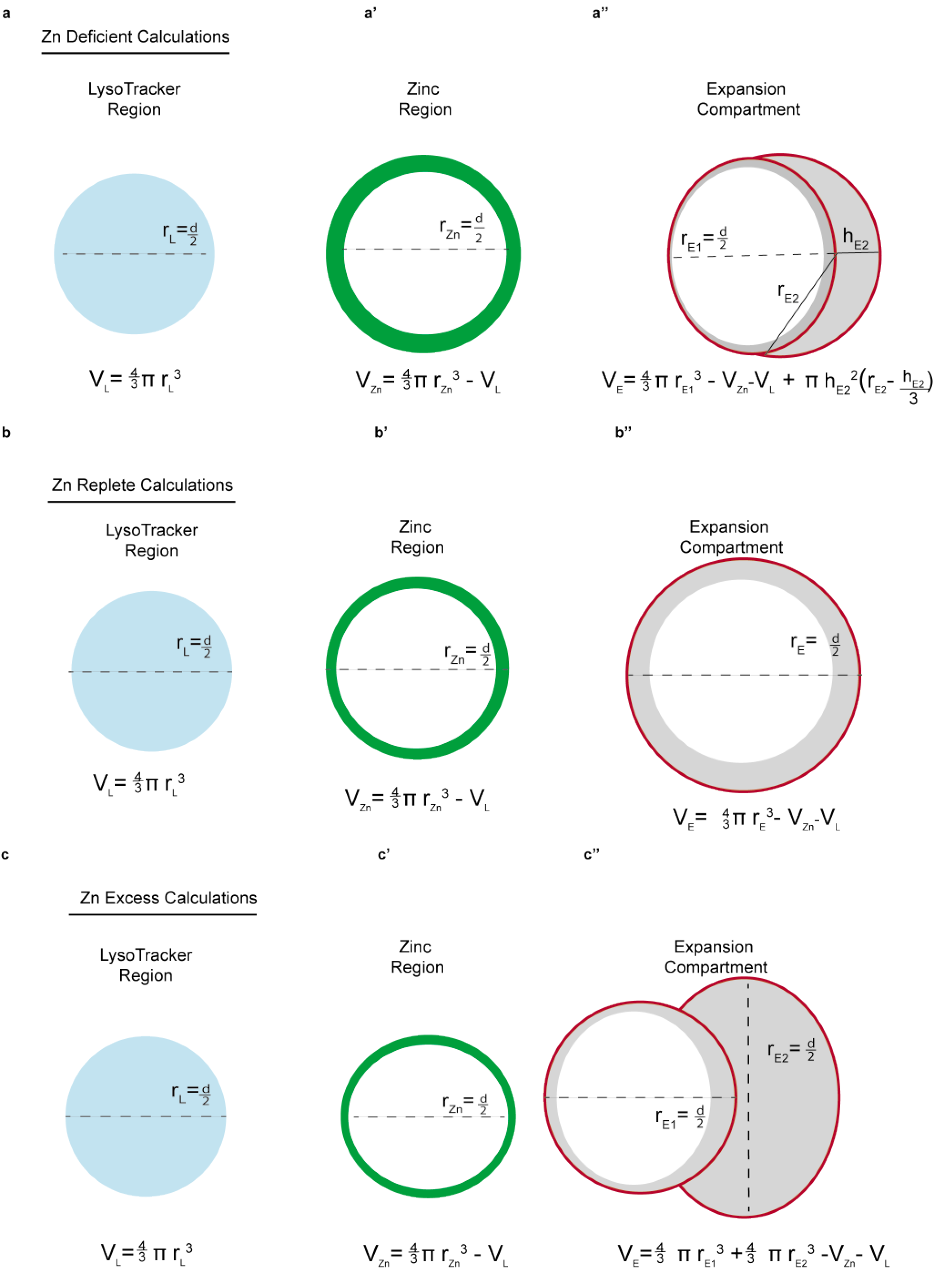
(with main Fig. 3e). Methods for calculating the volumes of the LysoTracker region, zinc region, and expansion compartment in zinc deficient, replete, and excess conditions. We developed mathematical equations to calculate the volumes of the LysoTracker region and the zinc region, which together comprise the acidified compartment, and the volume of the expansion compartment. (a-c) The LysoTracker region appears spherical in all conditions, and we used the same approach to calculate the volume (V_L_) in all conditions. We assumed it is a sphere, measured the diameter, calculated the radius (r_L_), and calculated the volume using the formula V_L_ = 4/3*π* r_L_^3^ (a’-c’). The zinc region appears to be a hollow sphere in all conditions, and we used the same approach to calculate the volume (V_Zn_) in all conditions. We assumed it is a sphere, measured the diameter, calculated the radius (r_Zn_), and calculated the volume using the formula V_Zn_ = 4/3*π*r_Zn_^3^, and then subtracted the volume of the LysoTracker region (V_L_). The sum of the volumes of the LysoTracker region and zinc region is the volume of the acidified compartment (V_A_ = V_L_ + V_Zn_) (a’’ – c’’). The expansion compartment displayed a distinct shape in each zinc condition, and we developed a unique formula for each condition. In zinc deficient conditions, we measured the expansion compartment volume by assuming it has two parts: (1) a hollow sphere surrounding the zinc region and (2) a hemispherical attachment. To calculate the total volume (V_E_), we independently measured the two parts. For part 1, we assumed it is a sphere, measured the diameter, calculated the radius (r_E1_), and calculated the volume using the formula V = 4/3 *π*r_E1_^3^, and then subtracted the volumes of the LysoTracker region (V_L_) and the zinc region (V_Z_). For part 2, we assumed it is a hemisphere, measured the radius (r_E2_) and the thickness (h_E2_), and used the formula, 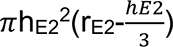. The total volume of the expansion compartment (V_E_) is the sum of the volumes of part 1 and part 2. In zinc replete conditions, we measured the expansion compartment volume by assuming it has one part - a hollow sphere surrounding the zinc region. We assumed it is a sphere, measured the diameter, calculated the radius (r_E1_), and calculated the volume using the formula V = 4/3 *π*r_E1_^3^, and then subtracted the volumes of the LysoTracker region (V_L_) and the zinc region (V_Z_). In zinc excess conditions, we measured the expansion compartment volume by assuming it has two parts: (1) a hollow sphere surrounding the zinc region and (2) a spherical attachment. To calculate the total volume (V_E_), we independently measured the two parts. For part 1, we assumed it is a sphere, measured the diameter, calculated the radius (r_E1_), and calculated the volume using the formula V = 4/3*π*r_E1_^3^, and then subtracted the volumes of the LysoTracker region (V_L_) and the zinc region (V_Z_). For part 2, we assumed it is a sphere, measured the diameter, calculated the radius (r_E2_), and calculated the volume using the formula V = 4/3*π*r_E2_^3^. The total volume of the expansion compartment (V_E_) is the sum of the volumes of part 1 and part 2.

**Supplemental Figure 6.**
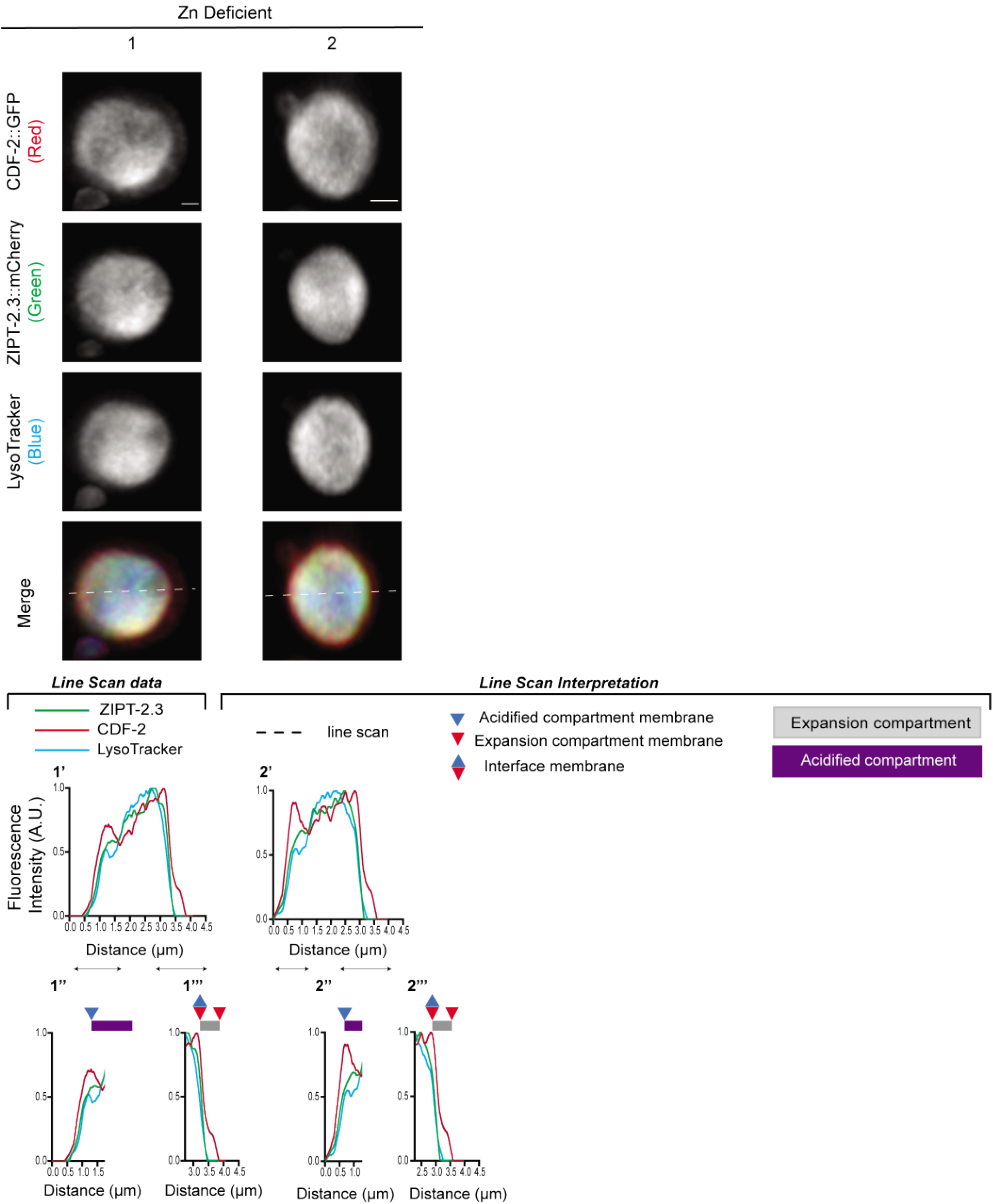
(with main Fig. 3a,b). Super resolution microscopy of gut granules in zinc deficient conditions with CDF-2, ZIPT-2.3 and LysoTracker. Transgenic L4 stage animals expressing CDF-2::GFP (true color green – arbitrary color red) and ZIPT-2.3::mCherry (true color red – arbitrary color green) were cultured for 16 hours in LysoTracker Blue (true color blue – arbitrary color blue) in 50 µM TPEN (Zn deficient). Individual gut granules were imaged by super resolution microscopy for green, red, and blue fluorescence, and a maximum intensity projection is displayed. Scale bar = 0.5 μm. (1’-2’) A line scan was performed, indicated by the dashed white line on merge image. For each color, the highest value was set equal to 1.0 arbitrary units (AU), and other values were normalized. (1’’-2’’) Enlargements of specific regions indicated by black lines. Annotations above indicate positions of membranes (triangles) and compartments (purple and gray rectangles).

**Supplemental Figure 7.**
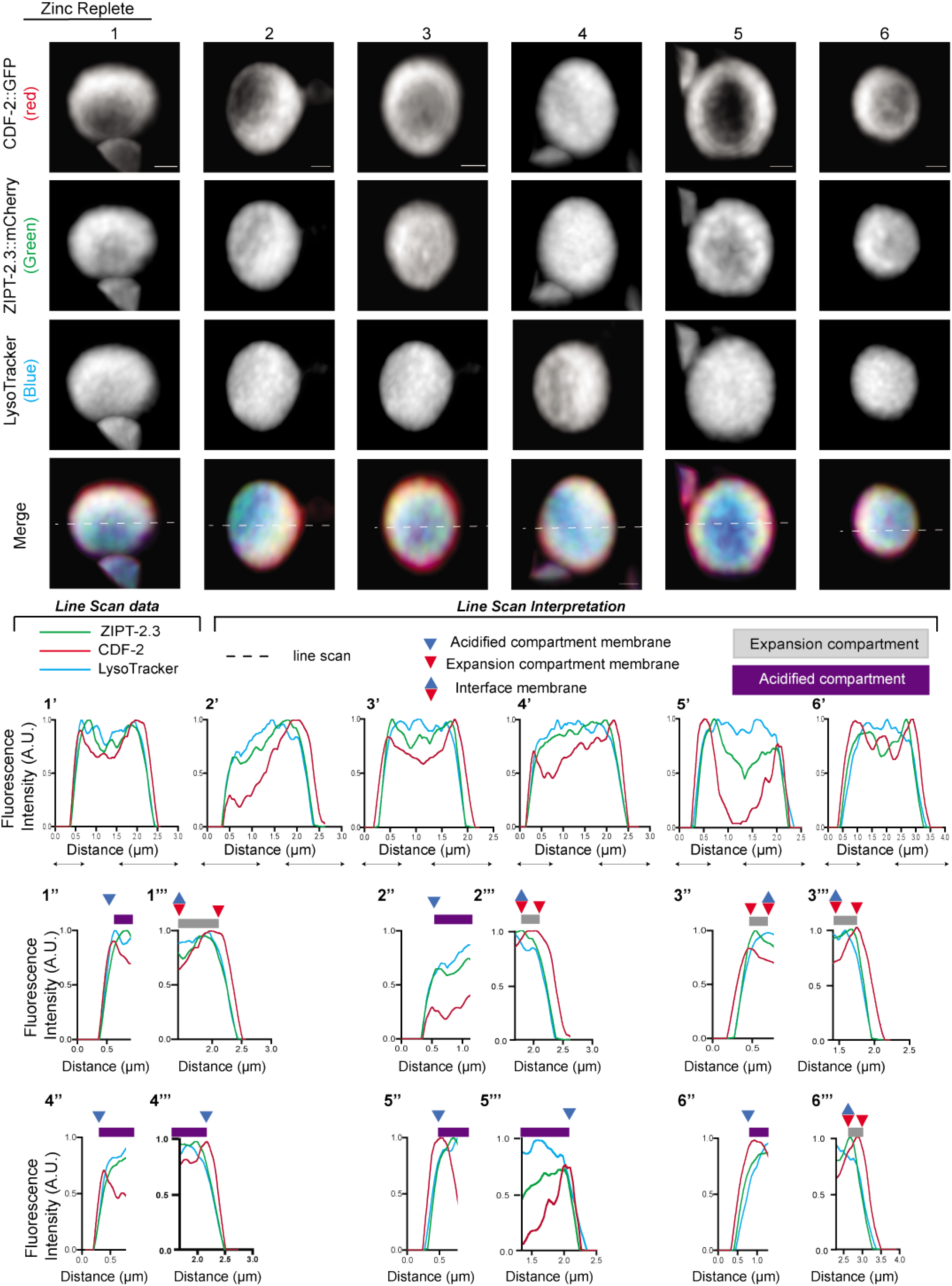
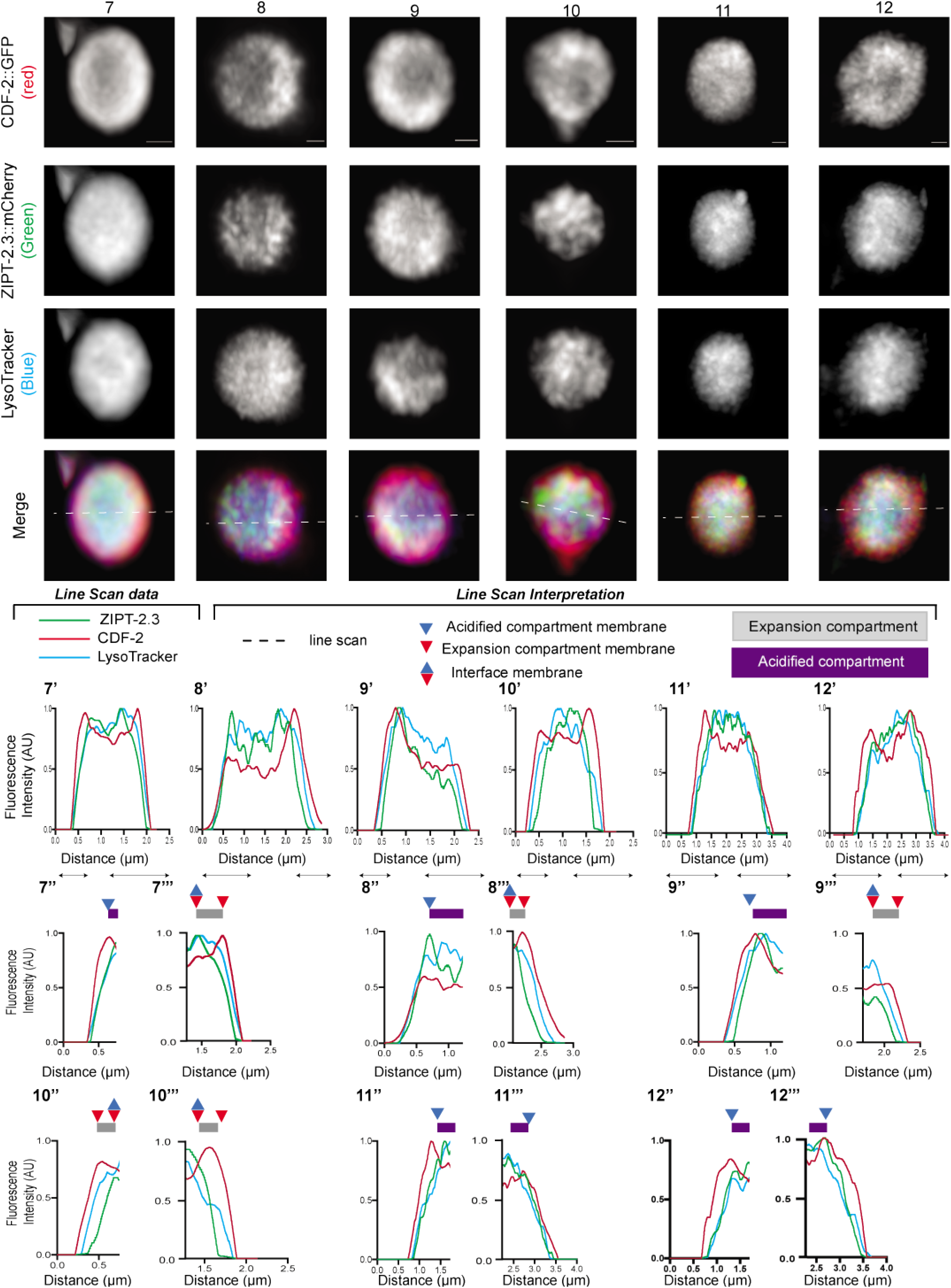
(with main Fig. 3a,c). Super resolution microscopy of gut granules in zinc replete conditions with CDF-2, ZIPT-2.3 and LysoTracker. Transgenic L4 stage animals expressing CDF-2::GFP (true color green – arbitrary color red) and ZIPT-2.3::mCherry (true color red – arbitrary color green) were cultured for 16 hours in LysoTracker Blue (true color blue – arbitrary color blue) in standard medium (Zn replete). Individual gut granules were imaged by super resolution microscopy for green, red, and blue fluorescence, and a maximum intensity projection is displayed. Scale bar = 0.5 μm. (1’-12’) A line scan was performed, indicated by the dashed white line on merge image. For each color, the highest value was set equal to 1.0 arbitrary units (AU), and other values were normalized. (1’’-12’’) Enlargements of specific regions indicated by black lines. Annotations above indicate positions of membranes (triangles) and compartments (purple and gray rectangles).

**Supplemental Figure 8.**
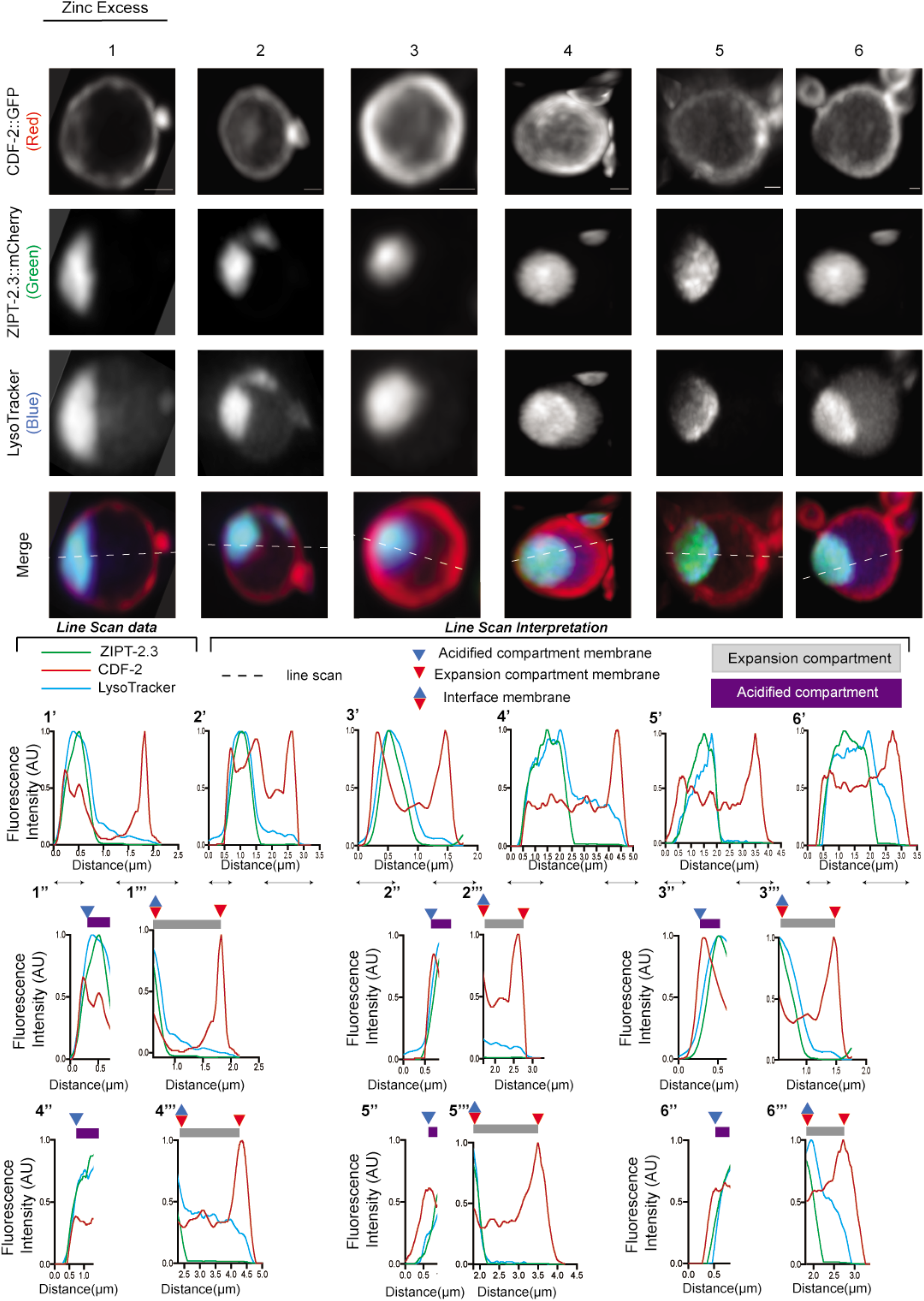
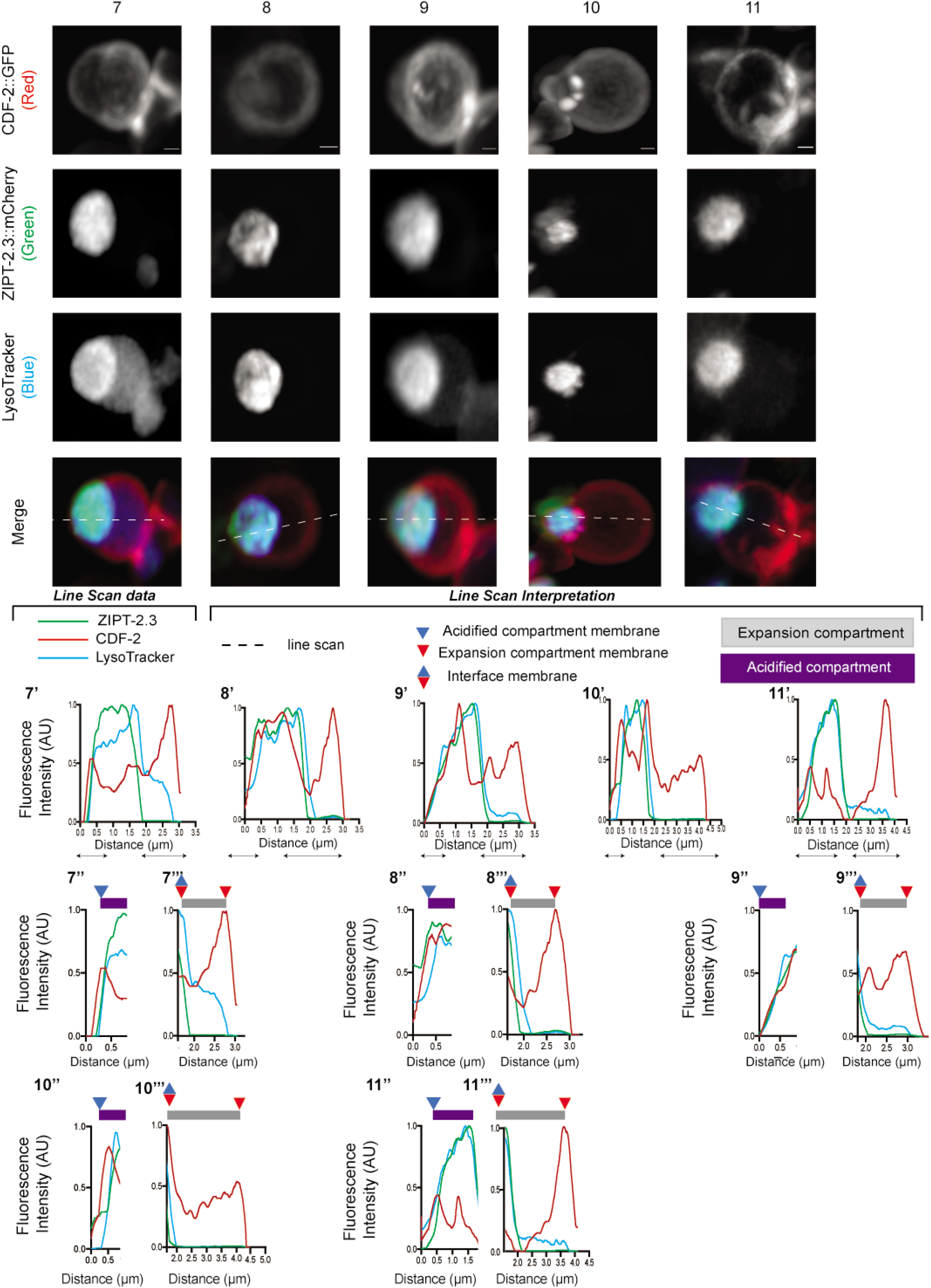
(with main Fig. 3a, d). Super resolution microscopy of gut granules in zinc excess conditions with CDF-2, ZIPT-2.3 and LysoTracker. Transgenic L4 stage animals expressing CDF-2::GFP (true color green – arbitrary color red) and ZIPT-2.3::mCherry (true color red – arbitrary color green) were cultured for 16 hours in LysoTracker Blue (true color blue – arbitrary color blue) in medium with 200 µM supplemental zinc (Zn excess). Individual gut granules were imaged by super resolution microscopy for green, red, and blue fluorescence, and a maximum intensity projection is displayed. Scale bar = 0.5 μm. (1’-11’) A line scan was performed, indicated by the dashed white line on merge image. For each color, the highest value was set equal to 1.0 arbitrary units (AU), and other values were normalized. (1’’-11’’) Enlargements of specific regions indicated by black lines. Annotations above indicate positions of membranes (triangles) and compartments (purple and gray rectangles).

**Supplemental Figure 9.**
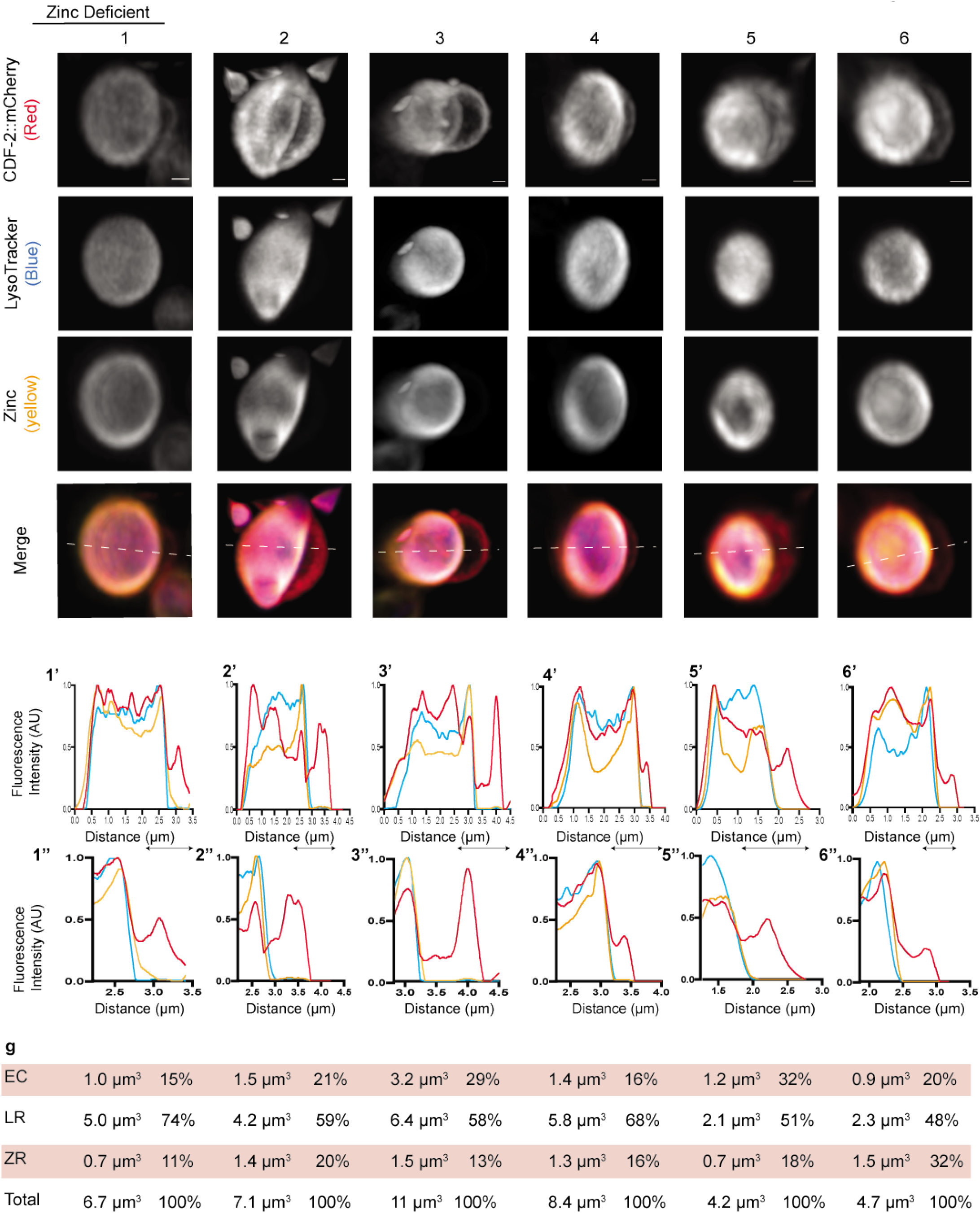
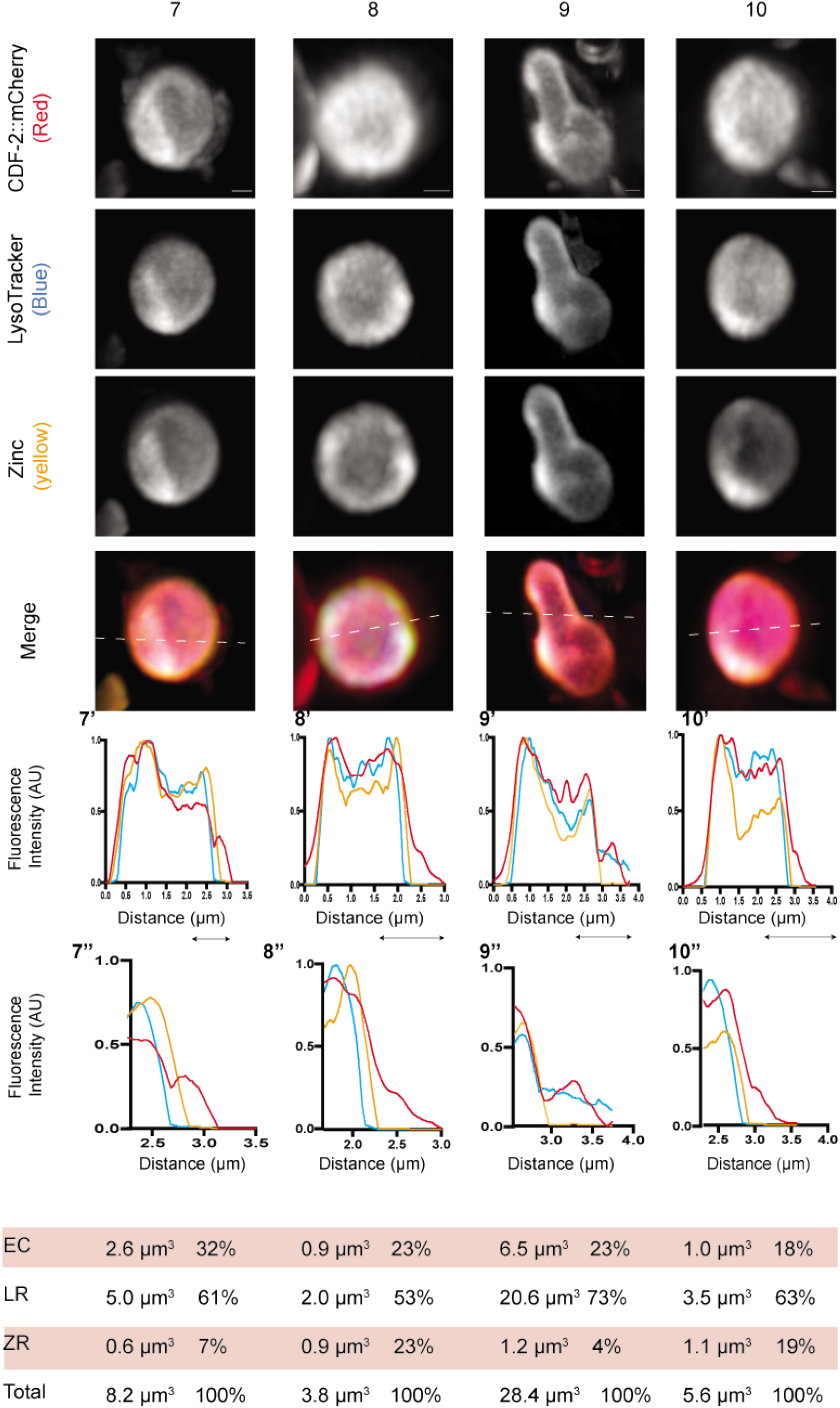
(with main Fig. 3e, f). Super resolution microscopy of gut granules in zinc deficient conditions with CDF-2, FluoZin-3 AM, and LysoTracker. Transgenic L4 stage animals expressing CDF-2::mCherry (true color red – arbitrary color red) were cultured for 16-20 hours in LysoTracker blue (true color blue – arbitrary color blue) and the zinc dye FluoZin-3 AM (true color green – arbitrary color yellow) in medium containing 50 µM TPEN (Zn deficient). Individual gut granules were imaged by super resolution microscopy for green, red, and blue fluorescence, and a maximum intensity projection is displayed. Scale bar = 0.5 μm. (1’-10’) A line scan was performed, indicated by the dashed white line on merge image. For each color, the highest value was set equal to 1.0 arbitrary units (AU), and other values were normalized. (1’’-10’’) Enlargements of specific regions indicated by black lines. Annotations above indicate positions of membranes (triangles), compartments (purple and gray rectangels) and regions (blue and orange rectangles). Gut granules were measured to determine the volume of the expansion compartment (EC), zinc region (ZR), and LysoTracker region (LR). Percent is the fraction of the total volume.

**Supplemental Figure 10.**
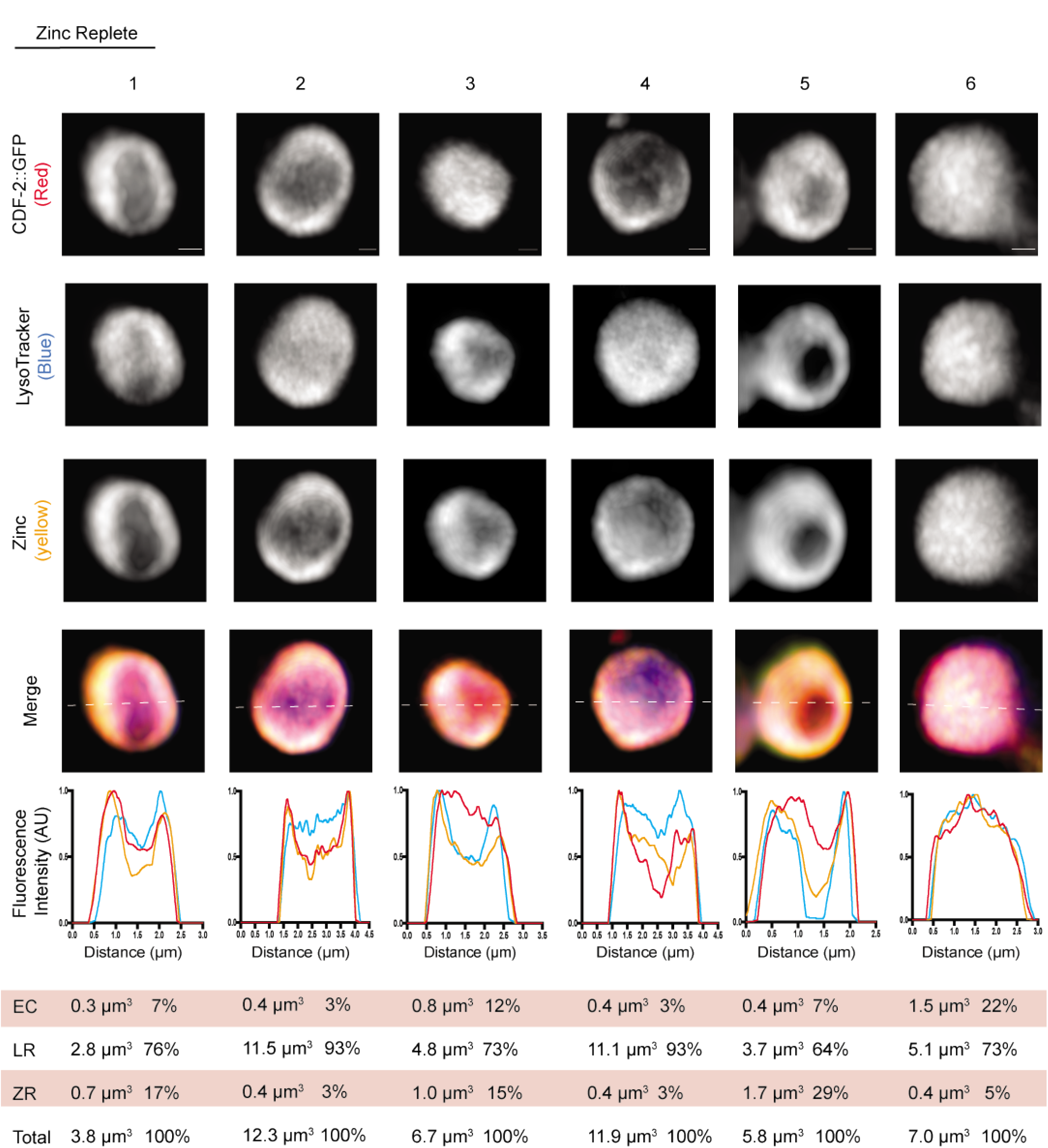
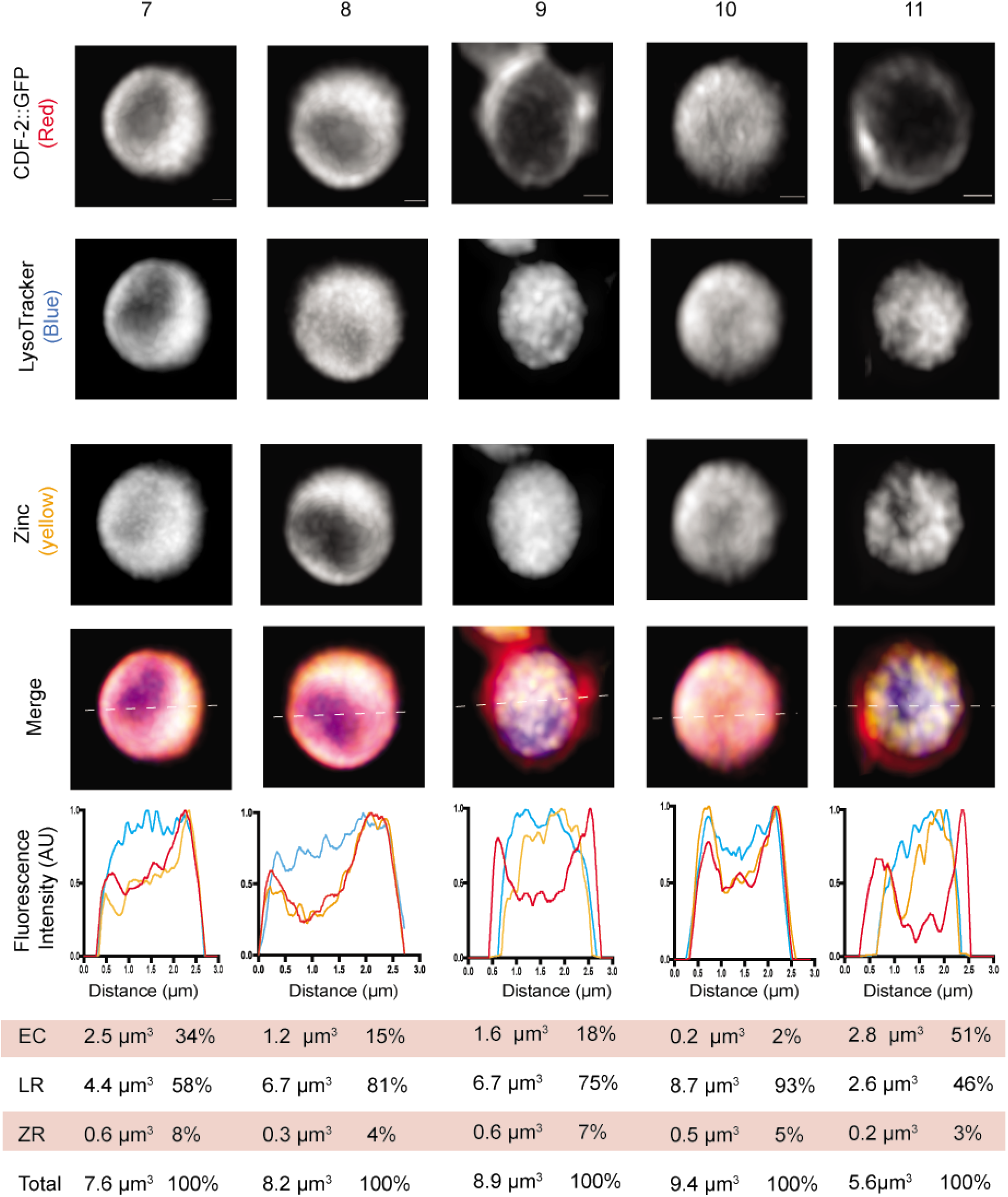
(with main Fig. 3e, g). Super resolution microscopy of gut granules in zinc replete conditions with CDF-2, FluoZin-3 AM, and LysoTracker. Transgenic L4 stage animals expressing CDF-2::mCherry (true color red – arbitrary color red) were cultured for 16-20 hours in LysoTracker blue (true color blue – arbitrary color blue) and the zinc dye FluoZin-3 AM (true color green – arbitrary color yellow) in standard medium (Zn replete). Individual gut granules were imaged by super resolution microscopy for green, red, and blue fluorescence, and a maximum intensity projection is displayed. Scale bar = 0.5 μm. (1’-11’)

**Supplemental Figure 11.**
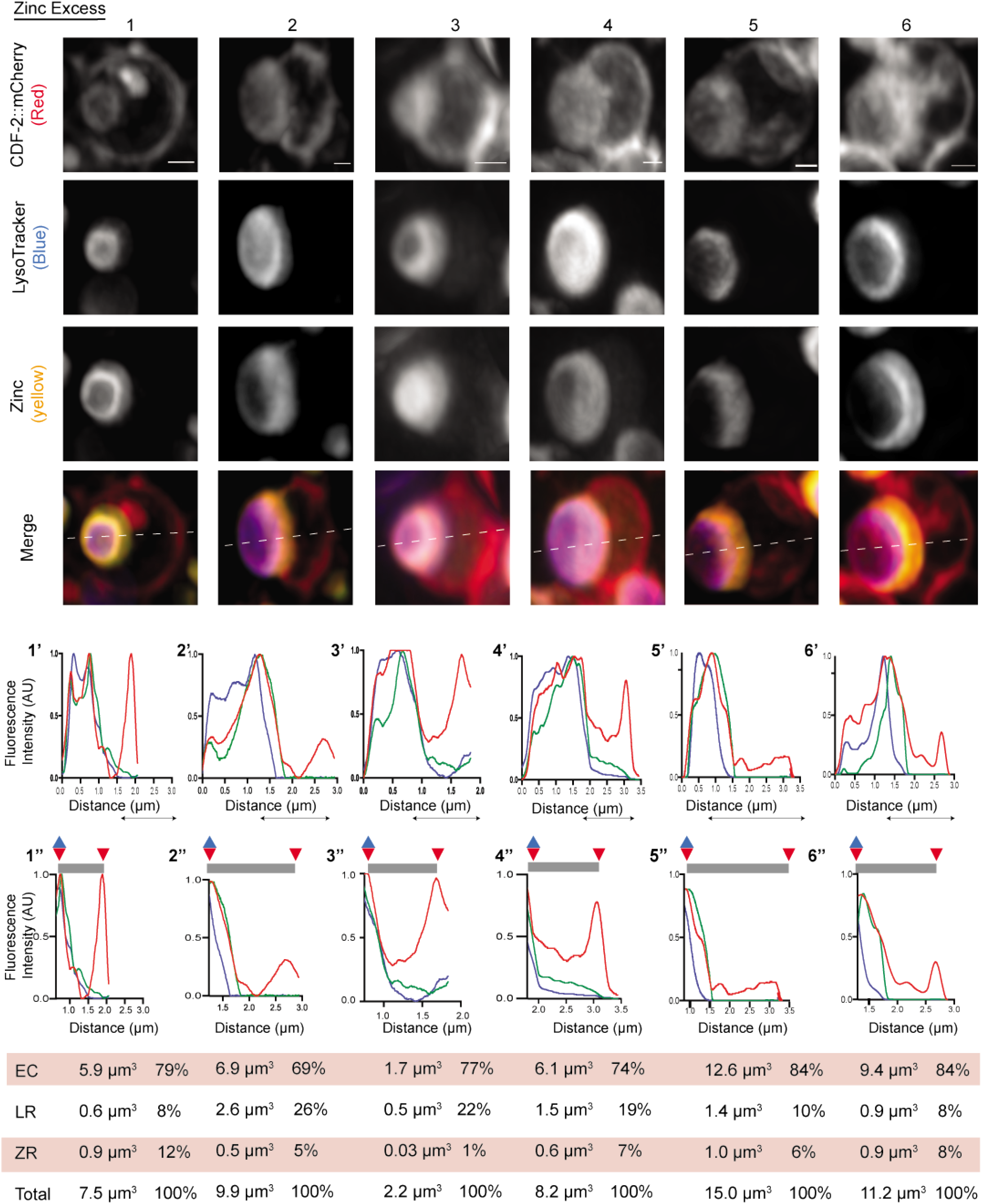
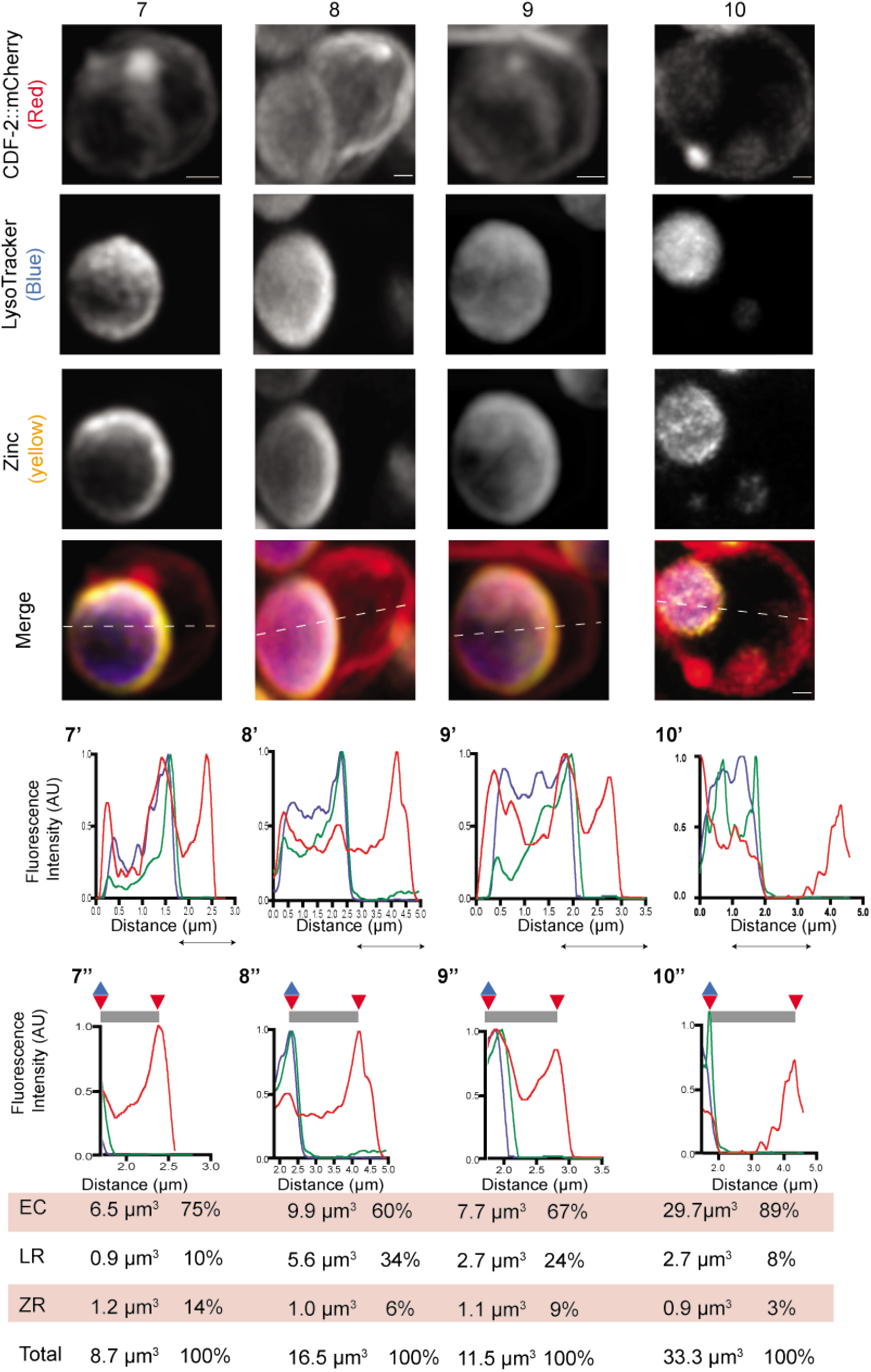
(with main Fig. 3eh). Super resolution microscopy of gut granules in zinc excess conditions with CDF-2, FluoZin-3, and LysoTracker. Transgenic L4 stage animals expressing CDF-2::mCherry (true color red – arbitrary color red) were cultured for 16-20 hours in LysoTracker blue (true color blue – arbitrary color blue) and the zinc dye FluoZin-3 AM (true color green – arbitrary color yellow) in medium containing 200 µM supplemental zinc (Zn excess). Individual gut granules were imaged by super resolution microscopy for green, red, and blue fluorescence, and a maximum intensity projection is displayed. Scale bar = 0.5 μm. (1’-11’) A line scan was performed, indicated by the dashed white line on merge image. For each color, the highest value was set equal to 1.0 arbitrary units (AU), and other values were normalized. Enlargements of specific regions indicated by black lines. Annotations above indicate positions of membranes (triangles) and compartments (purple and gray rectangels). Gut granules were measured to determine the volume of the expansion compartment (EC), zinc region (ZR), and LysoTracker region (LR). Percent is the fraction of the total volume.

**Supplemental Figure 12.**
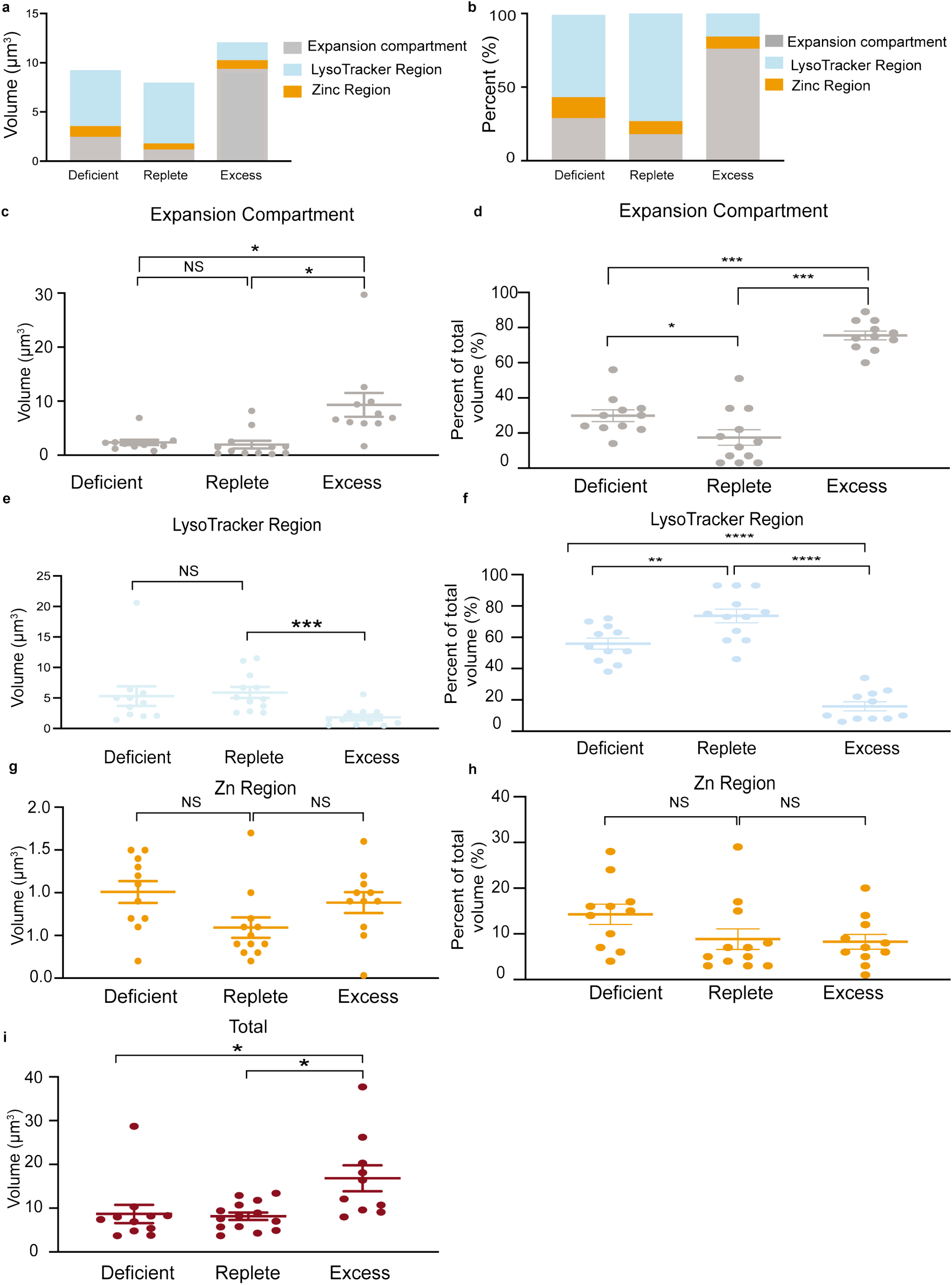
(with main Fig. 3e,k). Volumes of the LysoTracker region, zinc region, and expansion compartment in zinc deficient, replete, and excess conditions. Transgenic L4 stage animals expressing CDF-2::mCherry were cultured for 16-20 hours in LysoTracker blue and the zinc dye FluoZin-3 AM in either standard medium (Zn replete), 50 µM TPEN (Zn deficient) or 200 µM supplemental zinc (Zn excess). Individual gut granules were imaged by super resolution microscopy for green, red, and blue fluorescence. The volumes of the LysoTracker region, zinc region, and expansion compartment were measured (Fig. S5). (a-b) Bars represent the average volume (μm^3^) or the average percent of the total volume (%) of the zinc region (orange), Lysotracker region (blue), and expansion compartment (gray). N=11 deficient, 12 replete, and 11 excess. (c-i). Comparison of absolute volumes (μm^3^) or percentage of total volumes of the expansion compartment (c,d), LysoTracker region (e,f), zinc region (g,h), or total volume (i) in zinc deficient, replete, and excess conditions. Points are data from one organelle, and bar and whiskers indicate mean and standard error (* p<0.05, ** p<0.001, *** p<0.0001).

**Supplemental Figure 13.**
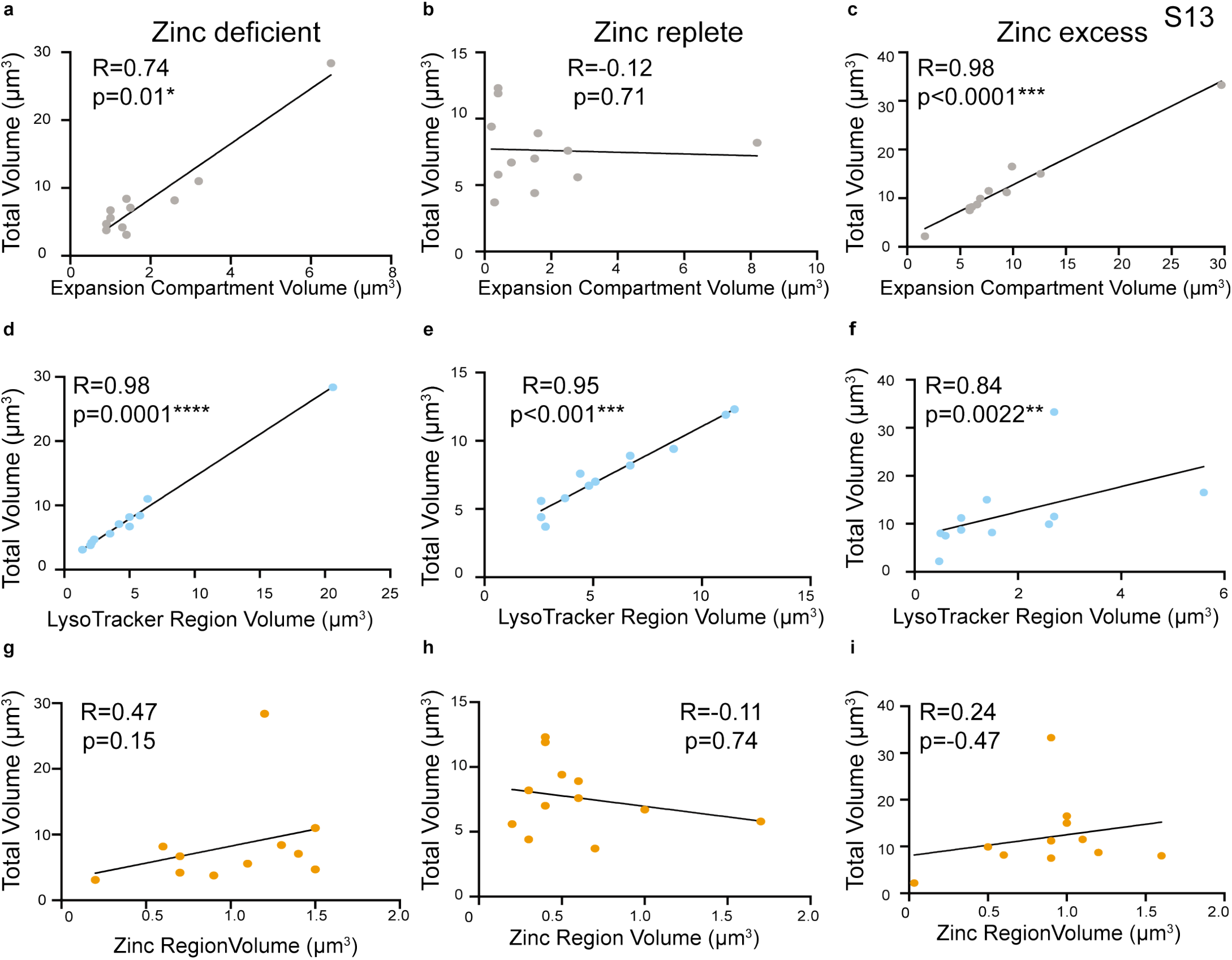
(with main Fig. 3e,k). Correlations between volumes of the LysoTracker region, zinc region, expansion compartment, and total volume in zinc deficient, replete, and excess conditions. Transgenic L4 stage animals expressing CDF-2::mCherry were cultured for 16-20 hours in LysoTracker blue and the zinc dye FluoZin-3 AM in either standard medium (Zn replete), 50 µM TPEN (Zn deficient) or 200 µM supplemental zinc (Zn excess). Individual gut granules were imaged by super resolution microscopy for green, red, and blue fluorescence. The volumes of the LysoTracker region, zinc region, and expansion compartment were measured (Fig. S5). (a-c) Data points represent the expansion compartment volume and the total volume of one gut granule. R is the correlation coefficient, where 1.0 is a perfect positive correlation, 0 is no correlation, and −1.0 is a perfect negative correlation. P is the likelihood that R is significantly different from 0 (* p<0.05, ** p<0.001, *** p<0.0001). There was a significant positive correlation in zinc excess and deficient conditions but not in zinc replete conditions. (d-f) Data points represent the LysoTracker region volume and the total volume of one gut granule. There was a significant positive correlation in all zinc conditions. (g-i) Data points represent the zinc region volume and the total volume of one gut granule. There was no significant correlation in any zinc condition.

## References

1. Andreini C, Banci L, Bertini I, Rosato A. Counting the zinc-proteins encoded in the human genome. J Proteome Res. 2006;5(1):196–201. Epub 2006/01/07. doi: 10.1021/pr050361j. PubMed PMID: 16396512.

2. Maret W. Zinc in Cellular Regulation: The Nature and Significance of “Zinc Signals”. Int J Mol Sci. 2017;18(11). Epub 2017/11/01. doi: 10.3390/ijms18112285. PubMed PMID: 29088067; PMCID: PMC5713255.

3. Qin Y, Dittmer PJ, Park JG, Jansen KB, Palmer AE. Measuring steady-state and dynamic endoplasmic reticulum and Golgi Zn2+ with genetically encoded sensors. Proc Natl Acad Sci U S A. 2011;108(18):7351–6. Epub 2011/04/20. doi: 10.1073/pnas.1015686108. PubMed PMID: 21502528; PMCID: PMC3088641.

4. Vinkenborg JL, Nicolson TJ, Bellomo EA, Koay MS, Rutter GA, Merkx M. Genetically encoded FRET sensors to monitor intracellular Zn2+ homeostasis. Nat Methods. 2009;6(10):737–40. Epub 2009/09/01. doi: 10.1038/nmeth.1368. PubMed PMID: 19718032; PMCID: PMC6101214.

5. Lichten LA, Cousins RJ. Mammalian zinc transporters: nutritional and physiologic regulation. Annu Rev Nutr. 2009;29:153–76. Epub 2009/04/30. doi: 10.1146/annurev-nutr-033009-083312. PubMed PMID: 19400752.

6. Kambe T, Tsuji T, Hashimoto A, Itsumura N. The Physiological, Biochemical, and Molecular Roles of Zinc Transporters in Zinc Homeostasis and Metabolism. Physiol Rev. 2015;95(3):749–84. Epub 2015/06/19. doi: 10.1152/physrev.00035.2014. PubMed PMID: 26084690.

7. Kimura T, Kambe T. The Functions of Metallothionein and ZIP and ZnT Transporters: An Overview and Perspective. Int J Mol Sci. 2016;17(3):336. Epub 2016/03/10. doi: 10.3390/ijms17030336. PubMed PMID: 26959009; PMCID: PMC4813198.

8. Conklin DS, McMaster JA, Culbertson MR, Kung C. COT1, a gene involved in cobalt accumulation in Saccharomyces cerevisiae. Mol Cell Biol. 1992;12(9):3678–88. Epub 1992/09/01. doi: 10.1128/mcb.12.9.3678. PubMed PMID: 1508175; PMCID: PMC360222.

9. MacDiarmid CW, Gaither LA, Eide D. Zinc transporters that regulate vacuolar zinc storage in Saccharomyces cerevisiae. EMBO J. 2000;19(12):2845–55. Epub 2000/06/17. doi: 10.1093/emboj/19.12.2845. PubMed PMID: 10856230; PMCID: PMC203372.

10. Docampo R, de Souza W, Miranda K, Rohloff P, Moreno SN. Acidocalcisomes - conserved from bacteria to man. Nat Rev Microbiol. 2005;3(3):251–61. Epub 2005/03/02. doi: 10.1038/nrmicro1097. PubMed PMID: 15738951.

11. Palmiter RD, Cole TB, Findley SD. ZnT-2, a mammalian protein that confers resistance to zinc by facilitating vesicular sequestration. EMBO J. 1996;15(8):1784–91. Epub 1996/04/15. PubMed PMID: 8617223; PMCID: PMC450094.

12. Roh HC, Collier S, Guthrie J, Robertson JD, Kornfeld K. Lysosome-related organelles in intestinal cells are a zinc storage site in C. elegans. Cell Metab. 2012;15(1):88–99. Epub 2012/01/10. doi: 10.1016/j.cmet.2011.12.003. PubMed PMID: 22225878; PMCID: PMC4026189.

13. Dietrich N, Schneider DL, Kornfeld K. A pathway for low zinc homeostasis that is conserved in animals and acts in parallel to the pathway for high zinc homeostasis. Nucleic Acids Res. 2017;45(20):11658–72. Epub 2017/10/05. doi: 10.1093/nar/gkx762. PubMed PMID: 28977437; PMCID: PMC5714235.

14. Zhao Y, Tan CH, Krauchunas A, Scharf A, Dietrich N, Warnhoff K, Yuan Z, Druzhinina M, Gu SG, Miao L, Singson A, Ellis RE, Kornfeld K. The zinc transporter ZIPT-7.1 regulates sperm activation in nematodes. PLoS Biol. 2018;16(6):e2005069. Epub 2018/06/08. doi: 10.1371/journal.pbio.2005069. PubMed PMID: 29879108; PMCID: PMC5991658.

15. Yin S, Qin Q, Zhou B. Functional studies of Drosophila zinc transporters reveal the mechanism for zinc excretion in Malpighian tubules. BMC Biol. 2017;15(1):12. Epub 2017/02/16. doi: 10.1186/s12915-017-0355-9. PubMed PMID: 28196538; PMCID: PMC5309981.

16. Qiu A, Shayeghi M, Hogstrand C. Molecular cloning and functional characterization of a high-affinity zinc importer (DrZIP1) from zebrafish (Danio rerio). Biochem J. 2005;388(Pt 3):745–54. Epub 2005/02/03. doi: 10.1042/BJ20041807. PubMed PMID: 15683366; PMCID: PMC1183453.

17. Dempski RE. The cation selectivity of the ZIP transporters. Curr Top Membr. 2012;69:221–45. Epub 2012/10/11. doi: 10.1016/B978-0-12-394390-3.00009-4. PubMed PMID: 23046653.

18. Chapman EM, Lant B, Ohashi Y, Yu B, Schertzberg M, Go C, Dogra D, Koskimaki J, Girard R, Li Y, Fraser AG, Awad IA, Abdelilah-Seyfried S, Gingras AC, Derry WB. A conserved CCM complex promotes apoptosis non-autonomously by regulating zinc homeostasis. Nat Commun. 2019;10(1):1791. Epub 2019/04/19. doi: 10.1038/s41467-019-09829-z. PubMed PMID: 30996251; PMCID: PMC6470173.

19. Davis DE, Roh HC, Deshmukh K, Bruinsma JJ, Schneider DL, Guthrie J, Robertson JD, Kornfeld K. The cation diffusion facilitator gene cdf-2 mediates zinc metabolism in Caenorhabditis elegans. Genetics. 2009;182(4):1015–33. Epub 2009/05/19. doi: 10.1534/genetics.109.103614. PubMed PMID: 19448268; PMCID: PMC2728845.

20. Moilanen LH, Fukushige T, Freedman JH. Regulation of metallothionein gene transcription. Identification of upstream regulatory elements and transcription factors responsible for cell-specific expression of the metallothionein genes from Caenorhabditis elegans. J Biol Chem. 1999;274(42):29655–65. Epub 1999/10/09. doi: 10.1074/jbc.274.42.29655. PubMed PMID: 10514435.

21. Hermann GJ, Schroeder LK, Hieb CA, Kershner AM, Rabbitts BM, Fonarev P, Grant BD, Priess JR. Genetic analysis of lysosomal trafficking in Caenorhabditis elegans. Mol Biol Cell. 2005;16(7):3273–88. Epub 2005/04/22. doi: 10.1091/mbc.e05-01-0060. PubMed PMID: 15843430; PMCID: PMC1165410.

22. Chazotte B. Labeling lysosomes in live cells with LysoTracker. Cold Spring Harb Protoc. 2011;2011(2):pdb prot5571. Epub 2011/02/03. doi: 10.1101/pdb.prot5571. PubMed PMID: 21285271.

23. Warnhoff K, Roh HC, Kocsisova Z, Tan CH, Morrison A, Croswell D, Schneider DL, Kornfeld K. The Nuclear Receptor HIZR-1 Uses Zinc as a Ligand to Mediate Homeostasis in Response to High Zinc. PLoS Biol. 2017;15(1):e2000094. Epub 2017/01/18. doi: 10.1371/journal.pbio.2000094. PubMed PMID: 28095401; PMCID: PMC5240932.

24. Aydemir TB, Liuzzi JP, McClellan S, Cousins RJ. Zinc transporter ZIP8 (SLC39A8) and zinc influence IFN-gamma expression in activated human T cells. J Leukoc Biol. 2009;86(2):337–48. Epub 2009/04/30. doi: 10.1189/jlb.1208759. PubMed PMID: 19401385; PMCID: PMC2726764.

25. Fukada T, Civic N, Furuichi T, Shimoda S, Mishima K, Higashiyama H, Idaira Y, Asada Y, Kitamura H, Yamasaki S, Hojyo S, Nakayama M, Ohara O, Koseki H, Dos Santos HG, Bonafe L, Ha-Vinh R, Zankl A, Unger S, Kraenzlin ME, Beckmann JS, Saito I, Rivolta C, Ikegawa S, Superti-Furga A, Hirano T. The zinc transporter SLC39A13/ZIP13 is required for connective tissue development; its involvement in BMP/TGF-beta signaling pathways. PLoS One. 2008;3(11):e3642. Epub 2008/11/06. doi: 10.1371/journal.pone.0003642. PubMed PMID: 18985159; PMCID: PMC2575416.

26. Jeong J, Walker JM, Wang F, Park JG, Palmer AE, Giunta C, Rohrbach M, Steinmann B, Eide DJ. Promotion of vesicular zinc efflux by ZIP13 and its implications for spondylocheiro dysplastic Ehlers-Danlos syndrome. Proc Natl Acad Sci U S A. 2012;109(51):E3530–8. Epub 2012/12/06. doi: 10.1073/pnas.1211775110. PubMed PMID: 23213233; PMCID: PMC3529093.

27. Amiri K, Kalish, A, Mukherji, S. Robustness and universality in organelle size control. bioRxiv. 2021;789453. doi: https://doi.org/10.1101/789453.

28. Brenner S. The genetics of Caenorhabditis elegans. Genetics. 1974;77(1):71–94. Epub 1974/05/01. PubMed PMID: 4366476; PMCID: PMC1213120.

29. Consortium CeDM. large-scale screening for targeted knockouts in the Caenorhabditis elegans genome. G3 (Bethesda). 2012;2(11):1415–25. Epub 2012/11/23. doi: 10.1534/g3.112.003830. PubMed PMID: 23173093; PMCID: PMC3484672.

30. Egan CR, Chung MA, Allen FL, Heschl MF, Van Buskirk CL, McGhee JD. A gut-to-pharynx/tail switch in embryonic expression of the Caenorhabditis elegans ges-1 gene centers on two GATA sequences. Dev Biol. 1995;170(2):397–419. Epub 1995/08/01. doi: 10.1006/dbio.1995.1225. PubMed PMID: 7649372.

31. Mello CC, Kramer JM, Stinchcomb D, Ambros V. Efficient gene transfer in C.elegans: extrachromosomal maintenance and integration of transforming sequences. EMBO J. 1991;10(12):3959–70. Epub 1991/12/01. PubMed PMID: 1935914; PMCID: PMC453137.

32. Schindelin J, Arganda-Carreras I, Frise E, Kaynig V, Longair M, Pietzsch T, Preibisch S, Rueden C, Saalfeld S, Schmid B, Tinevez JY, White DJ, Hartenstein V, Eliceiri K, Tomancak P, Cardona A. Fiji: an open-source platform for biological-image analysis. Nat Methods. 2012;9(7):676–82. Epub 2012/06/30. doi: 10.1038/nmeth.2019. PubMed PMID: 22743772; PMCID: PMC3855844.

